# easyCLIP Quantifies RNA-Protein Interactions and Characterizes Recurrent PCBP1 Mutations in Cancer

**DOI:** 10.1101/635888

**Authors:** Douglas F. Porter, Paul A. Khavari

## Abstract

RNA-protein interactions mediate a host of cellular processes, underscoring the need for methods to quantify their occurrence in living cells. RNA interaction frequencies for the average cellular protein are undefined, however, and there is no quantitative threshold to define a protein as an RNA-binding protein (RBP). Ultraviolet (UV) cross-linking immunoprecipitation (CLIP)-sequencing, an effective and widely used means of characterizing RNA-protein interactions, would particularly benefit from the capacity to quantitate the number of RNA cross-links per protein per cell. In addition, CLIP-seq methods are difficult, have high experimental failure rates and many ambiguous analytical decisions. To address these issues, the easyCLIP method was developed and used to quantify RNA-protein interactions for a panel of known RBPs as well as a spectrum of random non-RBP proteins. easyCLIP provides the advantages of good efficiency compared to current standards, a simple protocol with a very low failure rate, troubleshooting information that includes direct visualization of prepared libraries without amplification, and a new form of analysis. easyCLIP, which uses sequential on-bead ligation of 5’ and 3’ adapters tagged with different infrared dyes, classified non-RBPs as those with a per protein RNA cross-link rate of <0.1%, with most RBPs substantially above this threshold, including Rbfox1 (18%), hnRNPC (22%), CELF1 (11%), FBL (2%), and STAU1 (1%). easyCLIP with the PCBP1^L100^ RBP mutant recurrently seen in cancer quantified increased RNA binding compared to wild-type PCBP1 and suggested a potential mechanism for this RBP mutant in cancer. easyCLIP provides a simple, efficient and robust method to both obtain both traditional CLIP-seq information and to define actual RNA interaction frequencies for a given protein, enabling quantitative cross-RBP comparisons as well as insight into RBP mechanisms.

## Introduction

The number of RNA-protein interaction datasets is growing rapidly, raising the importance of being able to integrate them into models of the global RNA-protein interactome and the challenges of integrating such data between RNA-binding proteins (RBPs) was recently highlighted^1^. The physical reality of RNA-protein interactions is their individual occurrence in individual cells, which may be abstracted to an average complex number per-cell in a population. The RNA-protein complex count per-cell may be normalized to derive the number of complexes per-interaction partner. It is these frequencies, per-cell and per-interaction partner, that are the most basic characterizations of RNA-protein interaction networks. Determining the targets of an RBP by enrichment over negative control immunopurifications, or by clustering of cross-links, or many such other approaches, are all ultimately inferring that the absolute count of an RNA-protein complex in the cell is abnormally high. The estimation of per-cell and per-protein absolute quantities provide the ultimate framework for describing a global and widely reproducible view of RNA-protein interactions.

There is currently no general method to estimate absolute RNA-protein interaction frequencies, either by cross-linking or by other means. Relative interaction frequencies have been estimated by comparing co-purified radiolabeled RNA, but this method does not yield absolute numbers. It is possible to estimate cross-link rates by observing the amount of UV- and RNAse-dependent decrease in an immunoblot blot band for proteins that cross-link well, but this is not feasible for proteins with a cross-link rate of ~1%. Western blot quantification is further complicated by the fact that absolute quantification requires protein in single bands of at least 5 ng, the narrow region of linear signal in immunoblots, and the fact that protein cross-linked to an over-digested 1-3 base fragment of RNA (~0.3-1 kDa) will run so close to un-cross-linked protein that it would not be distinct for a ~70 kDa protein^2^.

One of the common questions in molecular biology is whether there are specific RNA interactions for a protein of interest, and what those RNAs are. However, there is no agreement on what constitutes a target RNA, and interactions occur along a continuum of affinities^3^. One potential criterion for a specific RNA interaction for a protein of interest is those interactions with a frequency per protein or fraction of interactions unlikely to occur with a randomly selected protein. Neither of these definitions have been used because no library of random non-RBP RNA-interactomes have been analyzed. One of the goals of this study was to enable target RNAs to be defined in these two ways.

Here we report an improvement to current CLIP protocols in an approach termed easyCLIP. easyCLIP reliably quantifies the numbers of RNA cross-links-per-protein and provides visual confirmation of each step in the CLIP protocol. easyCLIP was used to produce data for eleven randomly selected non-RBPs as well as a set of canonical RBPs, allowing us to approximate the distribution of RNA-binding interactions with the average protein and to propose a threshold for assignment of a protein as an RBP.

Finally, easyCLIP was applied to quantify the mutational impacts on RNA binding of L100 PCBP1. L100 PCBP1 missense mutants were highlighted in a recent global analysis of mutations in gastrointestinal adenocarcinoma (GIAC)^4^, which characterized a subset of GIAC that were broadly “genome stable” (i.e., lacking chromosome or microsatellite instability), but possessing frequent mutations in APC, KRAS, SOX9, and PCBP1. Unexpectedly, easyCLIP found the common cancer-associated L100 mutations in *PCBP1* increased the association of PCBP1 with RNA and suggested potential mechanisms for their selective advantage. easyCLIP is thus presented as a new CLIP method with built in verification checks that enables quantification of the number of RNA cross-links per protein to allow quantitative comparison across CLIP datasets.

## Results

### Library preparation by easyCLIP

To generate a simpler and faster way of producing CLIP-seq datasets, a method was developed using on-bead ligations^5–7^ of 3′ adapters (termed L3) and 5′ adapters (termed L5), each with a different fluorescent dye^8^ (Figure 1A, B). After running an SDS-PAGE gel and transferring to a nitrocellulose membrane, single- and dual-ligated RNA were clearly visible (Figure 1C). RNA was extracted from the nitrocellulose membrane using proteinase K, purified using oligonucleotide(dT) beads to capture the poly(A) sequence on the L3 adapter, eluted, reverse transcribed, and input directly into PCR (Figure 1A). Major differences from HITS-CLIP include the usage of a chimeric DNA-RNA hybrid for highly efficient ligation (see below), the purification of complexes from a gel by oligo(dT), L5 and L3 barcodes, UMIs, and the direct visualization of ligation efficiencies and finished libraries by infrared dyes (see below).

**Figure 1.**
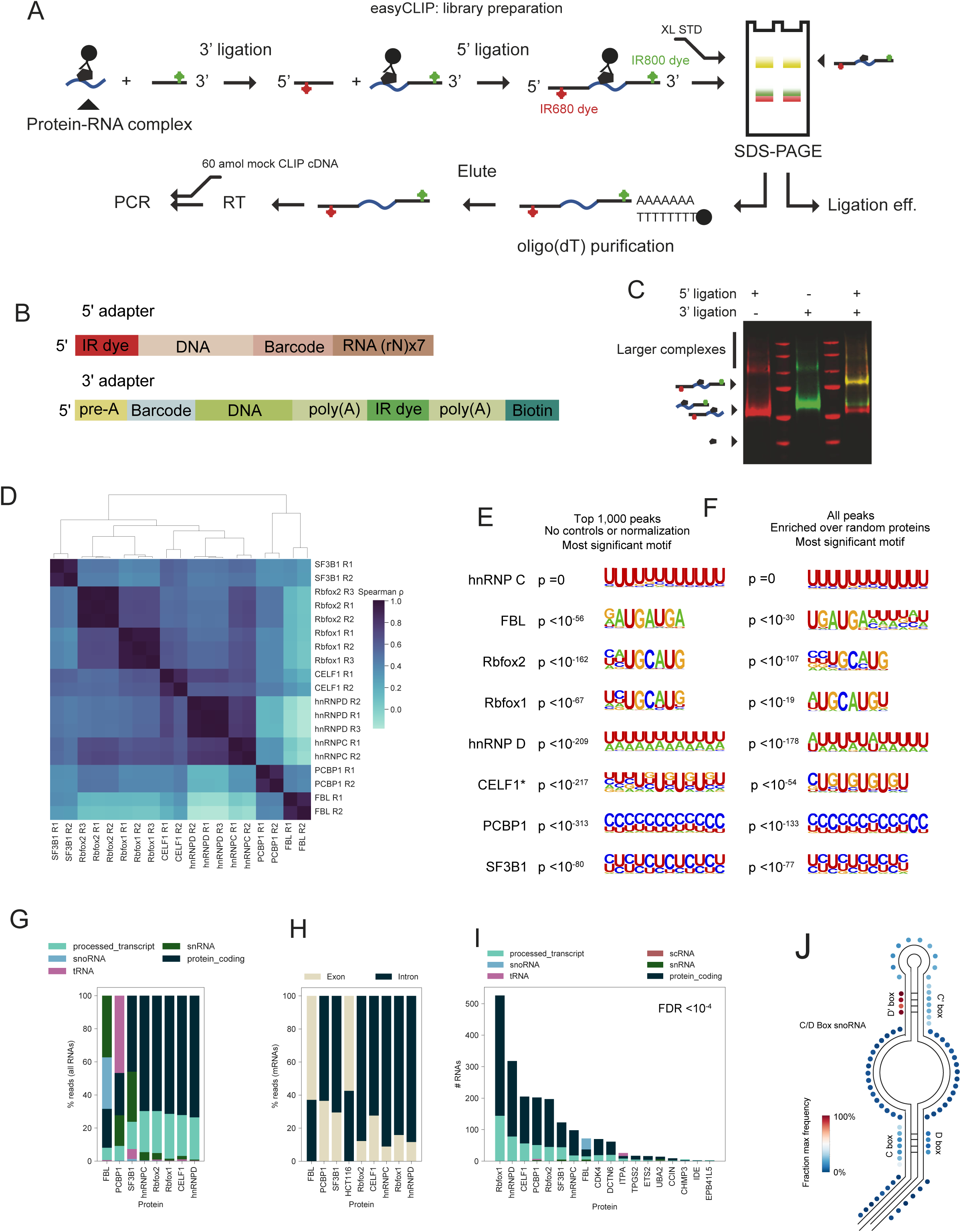
The easyCLIP protocol is a simpler and more quantitative CLIP-seq protocol. **A** The easyCLIP protocol to prepare libraries for high-throughput sequencing. **B** Diagrams of the 5′ and 3′ adapters used for the method in panel A and throughout this paper. Sequences are in File S1. The 5′ adapter has an IR680 dye (LI-COR) on blocking its 5′ end, and the 3′ adapter has an IR800-CW dye on an internal nucleotide. **C** A nitrocellulose membrane representing the gel in panel A, in which CLIP libraries produced for the protein hnRNP C are visualized by the dyes on the adapters, the 5′ in red and the 3′ in green. The left lane shows the products of only ligating the 5′ adapter, the middle lane only the 3′ adapter, and the right lane shows the products of ligating both the 5′ and 3′ adapters. The approximate size of hnRNP C (un-cross-linked) is indicated to the left. **D** Spearman ρ values for reads-per-gene from replicates of eight RBPs. **E** The most significant motif identified *de novo* by HOMER for the top 1,000 peaks. Specifically, the position of highest motif density in each RNA was identified by a method convolution of 5’ read positions across each RNA’s genomic locus and RNAs were ranked by their maximum height. The asterisk on CELF1 denotes that the motif was derived from the top 10,000 peaks. **F** The most significant motif identified *de novo* by HOMER for all peaks significantly enriched over randomly-chosen non-RBPs. Specifically, peak positions were identified as in panel E, but instead of taking the top 1,000, an FDR cutoff of 0.2 (Benjamini-Hochberg) was applied for enrichment of the given RNA (in reads-per-million) over the controls, and an FDR cutoff of 0.2 was applied for enrichment of reads-per-million at the given peak location over the controls. **G** The fraction of reads for easyCLIP of the indicated RBP that map to RNAs of the given ENSEMBL biotype. Labels are derived from the longest transcript isoform. The label *processed_transcript* indicates the RNA does not contain an ORF and is not given a more specific biotype (in this data, these reads largely map to lncRNA). Reads not mapping to one of the indicated categories are not counted in the total. **H** The fraction of reads in mRNA that map to introns or exons. **I** The number of significantly enriched (FDR<10^−4^) RNAs of the given category, relative to randomly selected non-RBP controls. **J** Fraction of FBL cross-link locations (identified as deletions) mapping to the given regions of a length-normalized C/D box snoRNA. Regions were normalized to their average length across all C/D box snoRNAs, and one dot represents one nucleotide in the normalized snoRNA. Frequencies are given as fractions of the nucleotide in the normalized snoRNA with the highest frequency.

This method (“easyCLIP”) incorporated several advantages. First, since all that happens after the gel extraction is a quick oligonucleotide(dT) purification and reverse transcription before PCR, there are minimal opportunities for error after the diagnostic step of gel imaging. Second, L5 and L3 barcodes may be used to mark samples and replicates, respectively, and all samples may be combined before running SDS-PAGE. This combination of samples allows for lower complexity preparations to “piggy-back” on higher complexity preparations, which allows very small RNA quantities that may be lost to sample absorption to be converted into libraries and for diagnostics from the larger libraries to be used for the smaller.

easyCLIP was benchmarked against eCLIP in the manner eCLIP was benchmarked against iCLIP, namely using Rbfox2, 10 µg antibody, and 20 million 293T cells. easyCLIP was more efficient than the published eCLIP results (Figure S1A), and easyCLIP RBFOX2 libraries fit expectations, including matching the pattern of binding seen with eCLIP at NDEL1 (Figure S1B), indicating that easyCLIP captures similar information. easyCLIP was then used to generate data for seven additional known RBPs: FBL (Fibrillarin, which associates with C/D-box snoRNA and other ncRNA), hnRNP C, hnRNP D, Rbfox1, CELF1, SF3B1 and PCBP1 (all of which at least partly bind mRNA). These were chosen as representatives (FBL, hnRNP C), for their importance to cancer (SF3B1, PCBP1), for comparison with eCLIP (Rbfox2), or by using a random number generator to select RBPs at random from the RBP atlas^9^ (Rbfox1, CELF1, hnRNP D). No randomly selected or representative RBPs were discarded.

easyCLIP libraries produced high quality data in each case (Figure 1D-J, Files 2-5). First, the data was consistent between replicates but distinct between proteins (Figure 1D). Second, FBL and hnRNP C/hnRNP D were un-correlated (Figure 1D), as expected. The data was high quality enough for all eight RBPs that simply feeding the sequences under the tallest 1,000 peaks (10,000 for CELF1) to a *de novo* motif discovery program^10^ resulted in the top motif being the expected motifs for all eight proteins, despite not performing any statistical tests, normalization, or comparison to a control (Figure 1E). This indicates easyCLIP data is clean enough that no statistical methods or controls are necessary to obtain good quality peaks. Using enrichment over controls also recovered all eight motifs (Figure 1F).

The motif obtained for FBL is expected because it is similar to the boxes of C/D box snoRNAs. As expected, hnRNP C, hnRNP D, CELF1, Rbfox1, and Rbfox2 bound mostly mRNA, while FBL was mostly crosslinked to snoRNA and snRNA (Figure 1G). PCBP1 and SF3B1 bound to both mRNA and snRNA, as expected. The main surprise was the appearance of tRNA-binding by PCBP1, addressed further below. About ~90% of hnRNP C/hnRNP D mRNA reads were intronic, as expected (Figure 1H). Under a highly stringent FDR<10^−4^ *vs* random non-RBPs (discussed below), target RNA numbers (Figure 1I) and the total number of unique mapped reads were both similar to what is typical for CLIP studies (Figure S1C); inputs ranged from a fraction of a 10 cm plate (Rbfox2, hnRNP C), to one 15 cm plate (PCBP1).

It is sometimes argued that iCLIP methods and their derivatives have higher resolution because the stop point of reverse transcriptase is mapped, but it has been shown that deletions in CLIP-seq reads also map binding sites to the same resolution^11^. For a very short RNA, such as a snoRNA, binding sites over much of the RNA are too close the 3’ end to be mappable, making the binding site ambiguous. However, using deletions allows binding sites anywhere in the RNA to be identified. Cross-linking positions within C/D box snoRNA were visualized in some detail (Figure 1J), and the respective frequencies of crosslinking in the different regions of C/D box snoRNAs matched previous reports^12^. This indicates easyCLIP provides an advantage over iCLIP/eCLIP-like methods for short RNAs, where reads with reverse transcriptase stops near the 3’ end are not mappable.

### Estimating absolute RNA quantities

easyCLIP was next tested to see if it could determine the total amount of RNA crosslinked to a given protein. Prior work has ligated 3′ adapter molecules labelled with infrared dyes to count crosslinked RNAs^8^, but this method does not account for un-ligated RNA, and is only accurate if there are no changes in dye fluorescence during the procedure or from imaging conditions.

When HEK293T cells were UV-crosslinked, hnRNP C immunopurified and RNA highly digested, a series of bands were visible by western blot (Figure 2A), spaced at roughly the ~60 kDa size of an hnRNP C dimer, as not all cross-linked complexes can be collapsed to monomers by RNAse digestion. If a ~15 kDa fluorescent adapter was ligated to highly digested hnRNP C-crosslinked RNA, a new band ~15 kDa above monomeric hnRNP C appeared containing adapter and hnRNP C (Figure 2B). The amount of protein in this band was determined by quantitative western blotting (Figure 2C, Figure S2A). The concentrations of standards were determined using multiple methods (Figure S2B-E), and consistency was established between epitope standards (Figure S2F). To determine if the hnRNP C antibody used (4F4) discriminated between non-cross-linked and cross-linked hnRNP C, epitope tagged hnRNP C was *in vitro* crosslinked to RNA, and 4F4 antibody showed only a negligible 16% bias (Figure S2G). Because the cross-linked band (Figure 2B) contains an equal number of protein and RNA molecules, quantification of the amount of protein in the cross-linked band relates adapter fluorescence values in this band into an absolute molecule number. Quantification of fluorescence per molecule using a single, large preparation of cross-linked, quantified hnRNP C as an aliquoted standard can be used to translate fluorescence values to RNA quantities if the loss is fluorescence is low and the ligation efficiency can be approximated.

**Figure 2.**
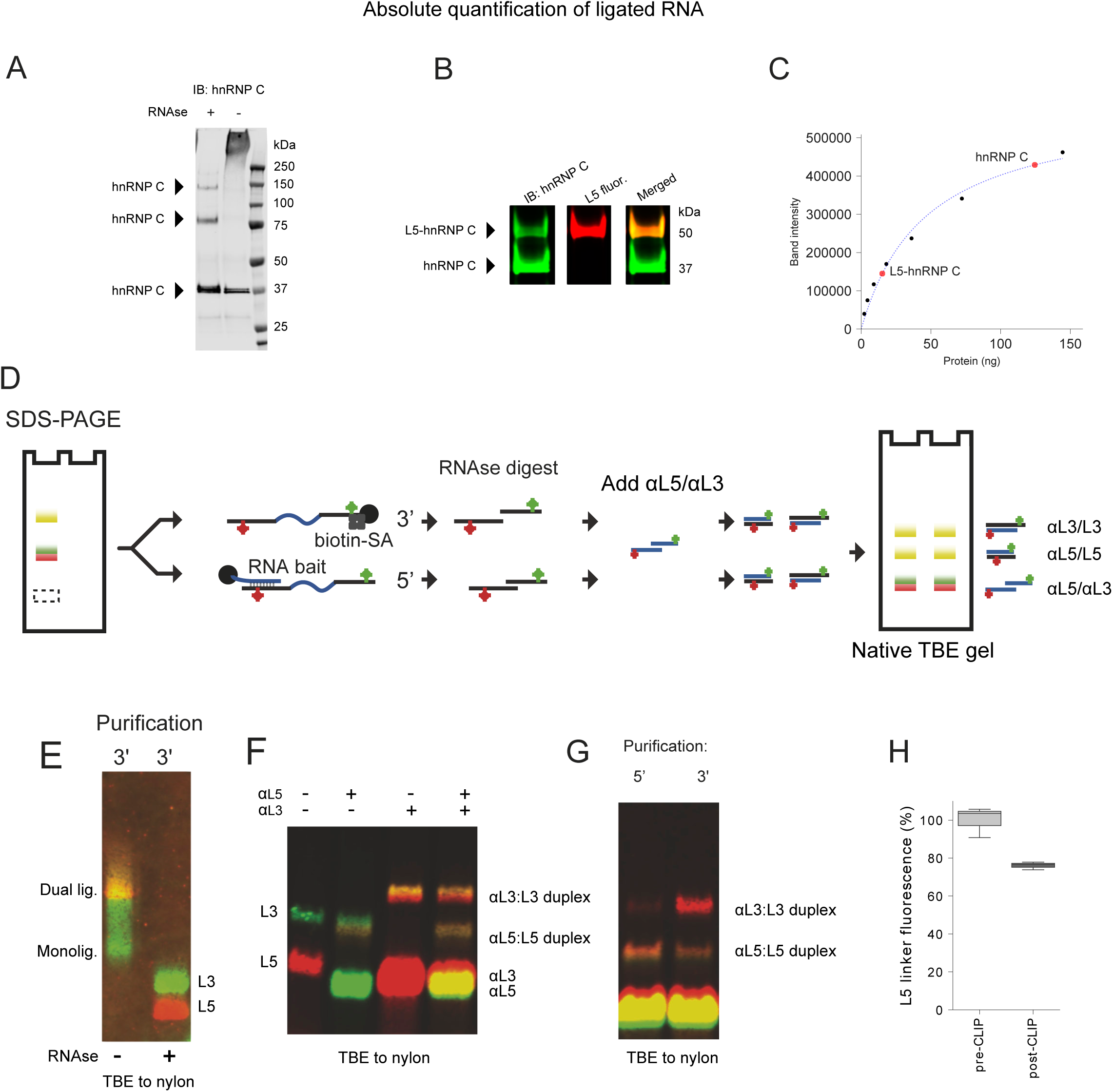
Development a method to quantify the absolute number of ligated RNA-protein complexes. **A** A large fraction of hnRNP C is UV cross-linked to RNA, much of which cannot be collapsed to a monomeric band by RNAse. **B** Ligation of the L5 adapter creates a novel band of monomeric hnRNPC-RNA-L5. Because this complex is the size of one hnRNP C molecule combined with one L5 molecule, the number of hnRNP C and L5 molecules in the L5-hnRNP C band are the same. **C** Absolute quantification of the monomeric hnRNP C and L5-hnRNP C band using a standard curve of GST-hnRNP C. This quantifies the number of L5 molecules in the complex and thereby fluorescence per molecule. **D** Diagram of a method to estimate the amount of fluorescence per L5 molecule lost in the CLIP procedure. RNA is extracted from nitrocellulose membranes by proteinase K digestion, the adapters are purified, RNAse digested, and paired with antisense oligonucleotides of the reciprocal color. The dsDNA pair of adapter and antisense oligonucleotide are run on a TBE gel, transferred to nylon and the ratio of adapter to antisense oligonucleotide is measured to determine the relative fluorescence of adapters having gone through CLIP *vs* fresh adapters. **E** RNA purified as in panel D may be RNAse treated to collapse signal into adapter bands. **F** The method in panel D applied to known amounts of free L5 and L3 adapter. Adapters are completely shifted. **G** Application of the method in panel D to visualize fluorescence of hnRNP C-crosslinked RNA after a CLIP procedure. **H** Fluorescence loss of L5 from the CLIP procedure is consistent between experiments at ~20%.

### Fluorescence loss

To address the loss in adapter fluorescence from CLIP, a method was developed to determine this value for labelled DNA oligonucleotides. Antisense oligonucleotides to L5 and L3 were labelled with reciprocal dyes, hereafter termed αL5 and αL3, and used to shift their cognate adapter. That is, a red αL3 and a green αL5 are used to shift a green L3 and red L5. Such antisense oligonucleotides shift the adapter molecules up in a native gel and produce bands of both colors with a 1:1 ratio of antisense and sense oligonucleotide (Figure 2D). L5 and L3 were successfully purified from proteinase K extract and RNAse digested down to free adapters (Figure 2E). 100% of L5 and L3 adapters were shifted in this manner (Figure 2F) and the method was applied to the RNAse digested CLIP oligonucleotides (Figure 2G). By comparing the ratio of αL5 to L5 for fresh L5 and L5 extracted from the nitrocellulose membrane in CLIP, the loss in L5 fluorescence from CLIP could be determined (Figure 2H). L5 consistently lost only ~20% of its fluorescence.

### Ligation efficiency

Three methods were used to estimate ligation efficiency. The most straightforward of these is to ligate both a fluorescent L5 and L3 adapter and visualize the single *vs* dual shift from one or both adapters being ligated (Figure 3A). By quantifying the amount of fluorescence signal in the single- and dual-ligated protein-RNA complexes, efficiency estimates are obtained for both 5′ and 3′ (Figure 3B and C). Assuming the two ligations are independent events, the total amount of crosslinked RNA is also obtained, including unlabeled RNA (Figure 3C). This method indicated that L5 ligation efficiencies were consistent and in the neighborhood of 50% (Figure 3D).

**Figure 3.**
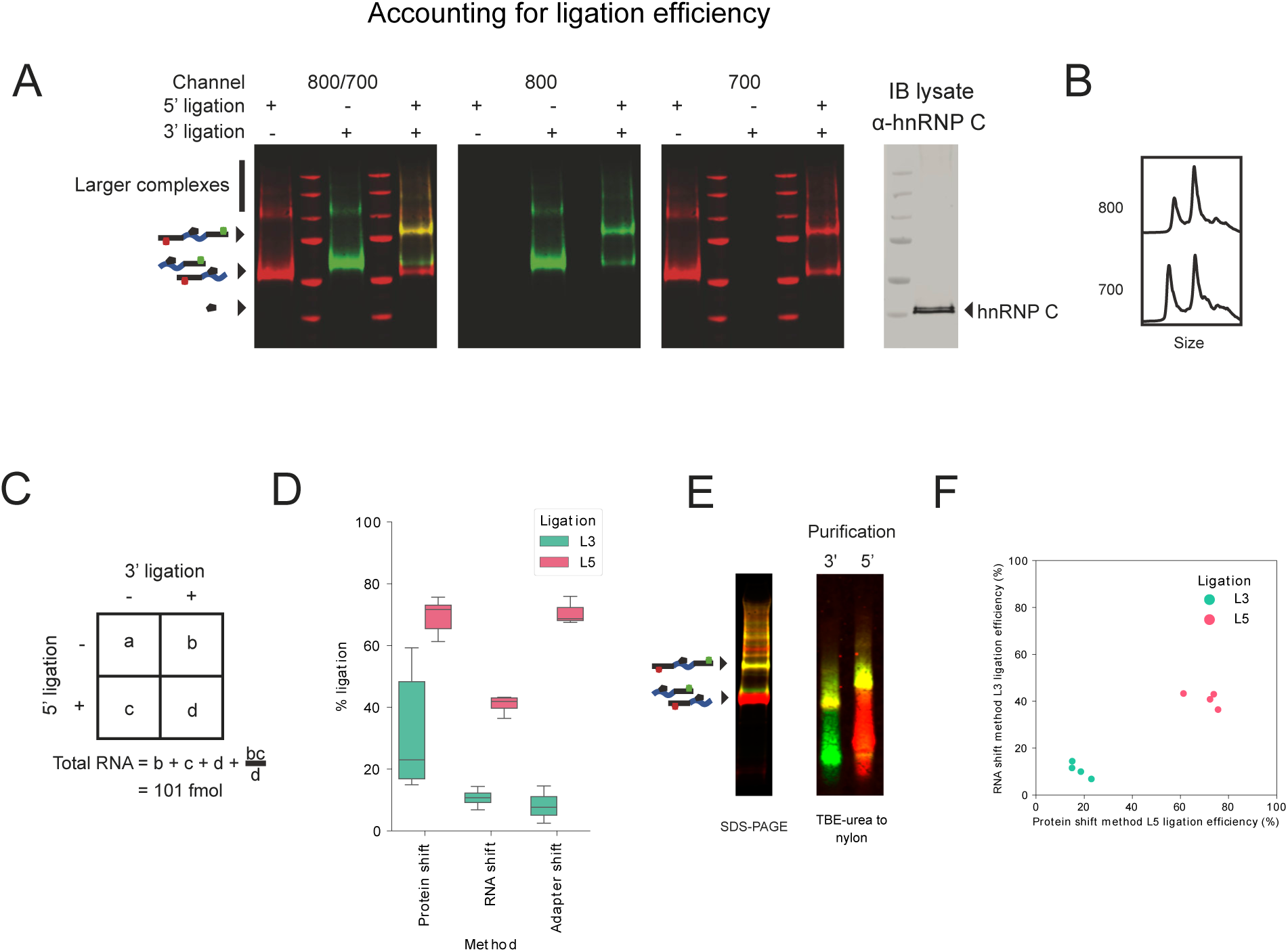
Estimating ligation efficiencies. **A** Products of single and dual ligations of the RNA-binding protein hnRNP C prepared according to the method in Figure 1A. Images represent nitrocellulose membranes, visualizing fluorescent markers ligated to RNA (left three panels) or a western blot of cell lysate (right panel). The observed ligation-mediated protein shifts form a method to determine to determine RNA ligation efficiencies in a CLIP protocol. **B** Chromatographs generated from the fluorescence gel image in panel A, far right lane. **C** Calculations to determine total bound RNA from the three observed values and the assumption of statistical independence in ligation efficiencies to determine the fourth value. **D** Ligation efficiencies estimated by the protein shift method (Figure 3A), by RNA shift (Figure 3E), or by shifting free adapters (Figure 2D). **E** Determination of ligation efficiency by shifted RNA. RNA was proteinase K extracted from nitrocellulose, adapters purified separately, intact RNA run on a gel and transferred to nylon. **F** Correlation between ligation efficiencies determined by the protein shift method or the RNA shift method for four biological replicates.

It was hypothesized that the higher molecular weight complexes visible in Figure 3A were produced by variation in the crosslinked protein, such as multimeric hnRNP C. If so, then the removal of protein by proteinase K digestion would remove the additional bands. To test this, RNA was extracted from nitrocellulose membranes using proteinase K, purified using either L5 or L3, run on a polyacrylamide gel, and transferred to a nylon membrane. Consistent with this hypothesis, higher molecular weight bands were collapsed into two simple smears of fluorescence, corresponding to mono-ligated and dual-ligated RNA (Figure 3E). A similar logic as applied in Figure 3A-C was applied to the protein-free RNA in Figure 3E to produce estimates of ligation efficiencies, which were lower but also consistent between replicates (Figure 3D, F).

A third method was also employed to quantify ligation efficiencies. Because the shifted bands in Figure 2G have a 1:1 ratio of L:αL oligonucleotides, quantifying antisense oligonucleotides also quantifies their respective adapters. The development of an antisense oligonucleotide-based method to quantify low femtomole amounts of adapter necessitated some optimization, described in Figures S3–8 and associated legends. For example, diluent has dramatic effects on fluorescence (Figure S5A) and there was a systematic test of the effects of salt, carrier, and PEG to retain fluorescence, prevent sample loss from adhesion, and preserve complexes on a gel (Figure S5B-F). Shifting known concentrations of L5 and L3 adapter fit well to a linear model, typically within 3 fmols (Figure S8C-D). By this third method, L5 ligation efficiencies were ~70% and consistent between CLIP rounds (Figure 3D). From these three methods, L5 ligation rates are stable between experiments and are roughly 50±20%. Altogether, results on the loss of adapter fluorescence (Figure 2) and ligation frequency (Figure 3) supported the use of standard aliquots (Figure 2B) to quantify absolute RNA amounts in CLIP experiments.

### Crosslink rates for RBPs

Two measures of RNA cross-linked to protein were determined: all RNA and minimal region RNA (Figure 4A). “All RNA” reflects the cross-linked RNA on the nitrocellulose membrane at the minimum size for a small protein-L5 complex (~30 kDa) and everything larger. Co-purified proteins cross-linked to RNA contribute to the total cross-linked RNA visualized. However, these are useful numbers because (1) since co-purified proteins must survive stringent purification conditions, they must constitute a close interaction of the protein of interest with RNA, and (2) the protein of interest often runs at a range of sizes (i.e. hnRNP C). The “minimal region” RNA measurement is taken from the region corresponding to the size for the dominant protein band cross-linked to small RNA fragments and ligated to L5, a region more likely to correspond to direct cross-linking events (Figure 4A). For all RNA, hnRNP C and FBL were 37% and 7% crosslinked to RNA, respectively (Figure 4B, see Figure S2H-I for FBL quantitative western blotting). Cross-link rates for the RBPs hnRNP D (19%), Rbfox1 (40%), CELF1 (21%), STAU1 (4.9%), PCBP1 (0.5%) and eIF4H (0.3%) were also established (Figure 4B). Cross-links in the minimal region (Figure 4C) were determined for RBPs hnRNP C (22%), FBL (2%), Rbfox1 (18%), CELF1 (11%) hnRNP D (5%), STAU1 (1.2%), PCBP1 (0.2%) and eIF4H (0.2%). STAU1 has a reputation as a very poor cross-linker^13^, so its cross-link rate may be taken as a representative for such.

**Figure 4.**
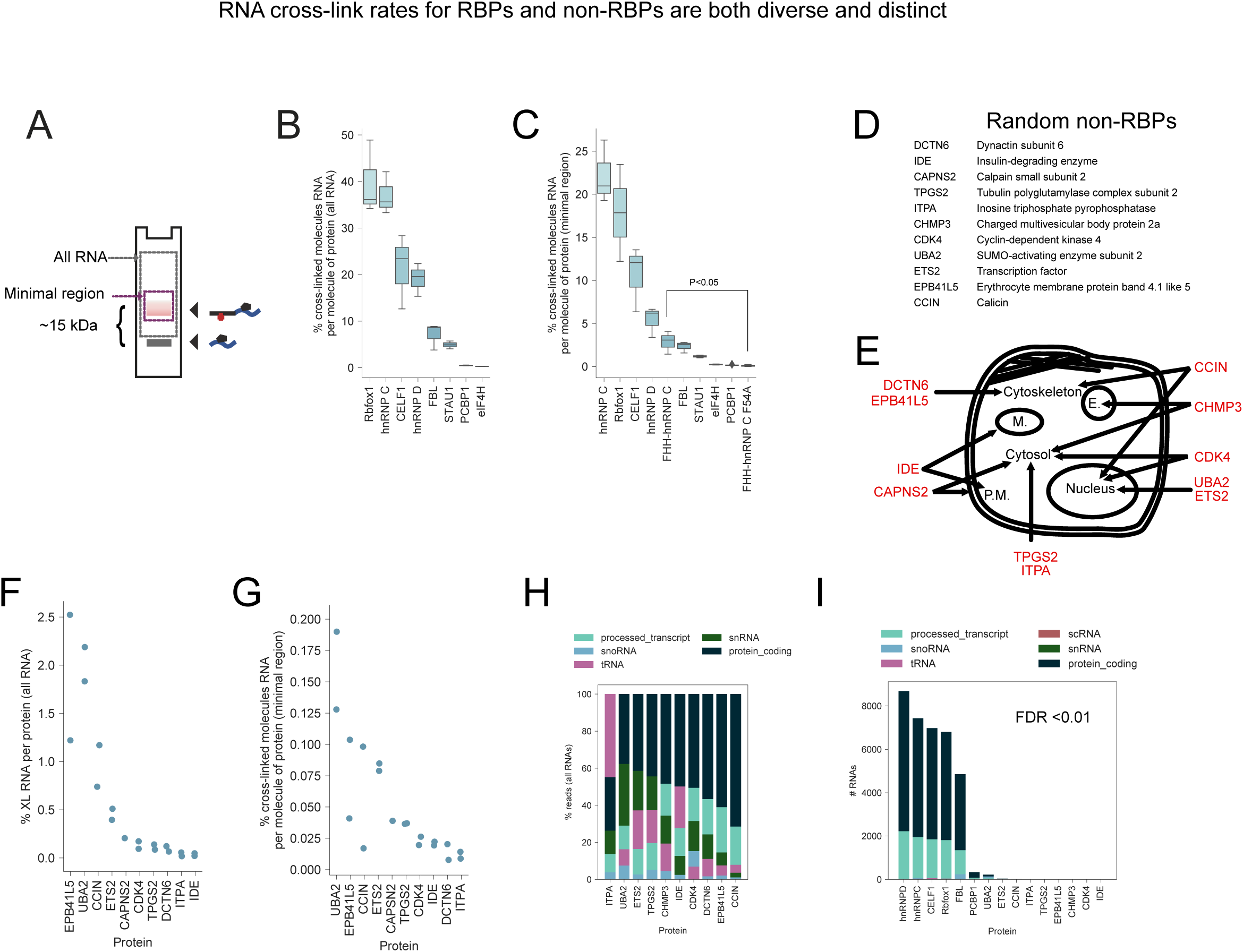
RNA cross-link rates for RBPs and non-RBPs are both diverse and distinct. Randomly selected non-RBPs provide a control for CLIP analysis. **A** Definition of the minimal region and all RNA/total RNA when quantifying the amount of purified, cross-linked RNA. **B** The protein cross-link rates of RBPs including all RNA. These rates were determined by quantitative western blotting for protein quantities, the determination of L5-ligated RNA by comparison of L5 fluorescence to a standard (Figure 2B), and the assumption of 50% ligation efficiency (Figure 3D). **C** The percent cross-linking to RNA for the indicated proteins, including only RNA within the minimal region. Tagged FHH-hnRNP C F54A could only be compared with FHH-hnRNP C by this method because both purify the wild-type hnRNP C, which is heavily cross-linked in either case. **D** Eleven randomly selected proteins that do not bind RNA *in vivo* or *in vitro*. **E** Subcellular locations of the randomly selected non-RBPs, as annotated on Uniprot. **F** UV cross-link rates to RNA for randomly selected non-RBPs, measuring all RNA. **G** UV cross-link rates to RNA for randomly selected non-RBPs, including only RNA in the minimal region. **H** The fraction of easyCLIP reads that map to RNAs of the given ENSEMBL biotype, excluding categories not shown. **I** The number of significantly enriched (FDR<0.01) RNAs of the given category for interactions *per protein* (*cf.* Figure 1I is *per read*), relative to randomly selected non-RBP controls.

The accuracy of this method was tested by calculating the cross-link rate of hnRNP C by quantitative western blotting of immunopurified hnRNP C (Figure S9). Results from this method agreed to within ~10%. It was asked if easyCLIP would reflect a loss in RNA-binding affinity caused by the F54A mutant of hnRNP C, a mutation in the RNA-binding surface of the RRM that elevates the RRM’s *in vitro* K_D_ from ~1 µM to >20 µM^14^. The mutant was dramatically less cross-linked (Figure 4C, P<0.05 t-test), although it still cross-linked better than the average human non-RBP (discussed below), consistent with hnRNP C functioning in a complex and possessing RNA contacts outside the RRM.

### Cross-link rates for non-RBPs

Quantification of cross-link rates may identify a numerical threshold for distinguishing RBPs from non-RBPs and for determining when an RBP has lost or gained RNA-binding activity. To derive a distribution of cross-link rates for non-RBPs, 11 non-RBP proteins were randomly selected using a script. This set of randomly selected non-RBPs cover a diverse range of functions and subcellular locations (Figure 4C-D). Selected non-RBPs had total RNA crosslink values of 0.03-2% (Figure 4F-G, Figure S10), and rates correlated with protein size (Figure S10B). Reducing counts to minimal region RNA dropped all cross-link rates except UBA2 to below 0.1%, and UBA2 to 0.16% (Figure 4G).

### Cross-linking rates distinguish RBPs and non-RBPs

Data above indicate that cross-link rates derived from a minimal region are typically below 0.1% for non-RBPs and above 0.1% for RBPs. The amount of total cross-linked RNA purified, not just that in the minimal region, ranges greatly (non-RBPs 0.1-2%, RBPs 0.2-42%). These metrics can be used to aid in defining what proteins are RBPs. For example, FHH-hnRNP C F54A had a minimal region cross-link rate of 0.1%, consistent with losing most direct affinity for RNA but still joining an RNA-binding complex.

### Defining specific interactions of RBPs and non-RBPs

One of the goals of this study was to enable target RNAs to be defined for a protein of interest as those interactions with a frequency per protein or per-cross-link unlikely to occur with a randomly selected protein. To do so, easyCLIP libraries for the ten of the eleven random non-RBPs were prepared. The specificity of the resulting libraries was confirmed by the over-representation of each overexpressed protein’s own RNA in CLIP data (Figure S10C). Despite not being RBPs, different non-RBPs produced distinct RNA-interactions (Figure 4H, Figure S10D). The two solely-nuclear proteins UBA2 and ETS2 had a low fraction of mRNA reads (Figure 4H).

Using the resulting distribution of RNA interactions for random proteins, it is possible to directly estimate how “unusual” any RNA-protein interaction pair is. This method was first applied to interaction frequencies per cross-link (i.e. per read). The validity of this method is supported by the identification of the expected motif for all eight RBPs as the top motif (Figure 1F), and target RNA types were consistent with expectations (Figure 1I): FBL targeted snoRNA, while hnRNPs targeted mRNA, and the core snRNP component SF3B1 targeted mRNA and snRNA. The number of FBL mRNA targets at least partly reflects mRNAs containing intronic snoRNAs. For each non-RBP, its own targets were defined after removing it from the set of controls, yet this still resulted in few “target” RNAs.

Finally, we identified target RNAs as those bound *per protein* at an unusually high rate. Frequent mRNA and lncRNA interactions per protein are characteristic of RBPs (Figure 4H). The rate of cross-linking per protein was plotted as a histogram to all mRNAs (Figure 5, *left*), snoRNAs (*middle*), or tRNA (*right*), which suggested some fundamental results. First, the distribution of binding across mRNAs, in reads-per-million, is similar between RBPs and non-RBPs (*top left*), but RBPs have many more frequent mRNA partners per protein. snoRNA presents a different picture (*middle*). Naïvely, if one looked only at reads-per-million, it would seem that either randomly selected proteins target snoRNA, or else RBPs somehow specifically avoid it. Per-protein, however, mRNA-binding RBPs and non-RBPs are equally likely to contact snoRNA – consistent with only FBL having specific interactions with snoRNA (*bottom middle*). The reason for this is clear enough – mRNA-binding RBPs have additional interactions that decrease the fraction of total interactions that occur with snoRNA. Despite its extremely high cross-link rate to mRNA, hnRNP C cross-links to snoRNA the same a random protein, as expected from such interactions being random. This cautionary tale helps explain the tRNA-binding observed by PCBP1 (Figure 1G). Like snoRNAs, tRNAs make up a disproportionate share of the libraries of non-RBPs (*top right*), but per-protein all RBPs and non-RBPs have the same distribution (*bottom right*). The distribution of tRNA binding by PCBP1 is actually just that of a non-RBP, indicating that it has no evolved interaction with tRNA, as might have been thought from conventional analysis in the absence of randomly selected non-RBPs.

**Figure 5.**
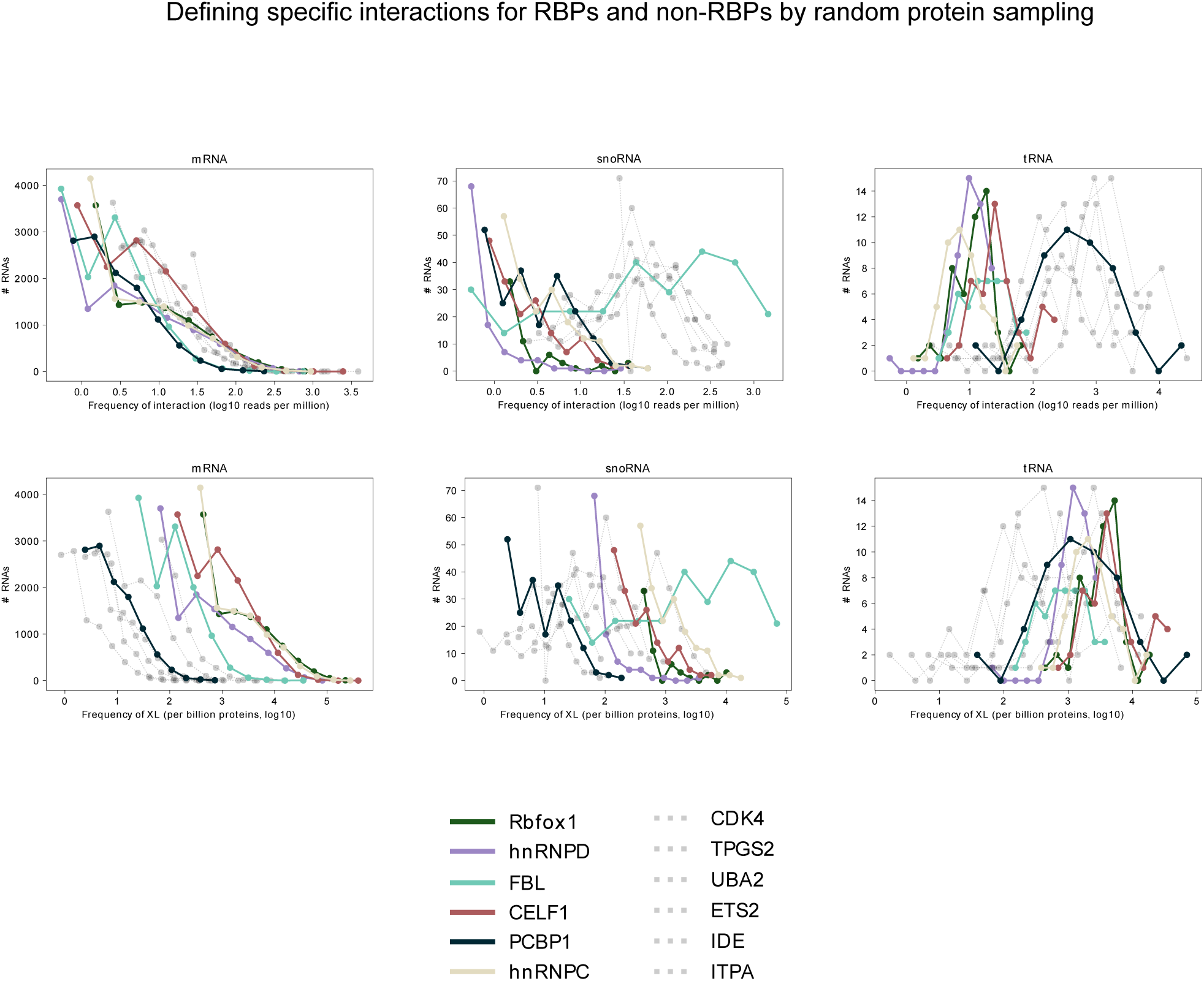
easyCLIP enables a new form of CLIP analysis that clarifies *in vivo* RNA-protein interaction landscapes. This figure depicts histograms (points indicate the left edge of each histogram bin, lines are for easier visualization) of cross-links per protein for RBPs and randomly selected non-RBPs. *Top* The x-axis is the rate of cross-linking per million cross-links molecules (i.e., reads per million) to a given RNA converted to a log10 scale, and y-axis is the number of RNAs at that cross-link rate. For each protein, RNAs with no reads were removed before determining the histogram (hence the leftmost bin varies by dataset size). *Bottom* The x-axis is the rate of cross-linking per billion protein molecules to the given RNA (i.e., reads per million, per million proteins), converted to a log10 scale, and the y-axis is the number of RNAs at that cross-link rate. RNAs with no reads were not included in the histogram. For both *top* and *bottom*, RNAs that would be placed outside beyond the rightmost bin were placed in the rightmost bin. In all three categories, the visualization *by protein* corresponds more to the expected biological reality.

### Cancer-associated mutations

The most frequent missense mutations in RBPs were identified in cancer using TCGA data^15^ (Figure 6A). The K700E mutant of SF3B1, the L100P/L100Q mutants of PCBP1 (Figure 6B), and the P131L mutant of RQCD1 were selected. SF3B1 K700E and RQCD1 P131L did not have obvious effects on RNA-binding in preliminary experiments, so PCBP1 was focused on for analysis.

**Figure 6.**
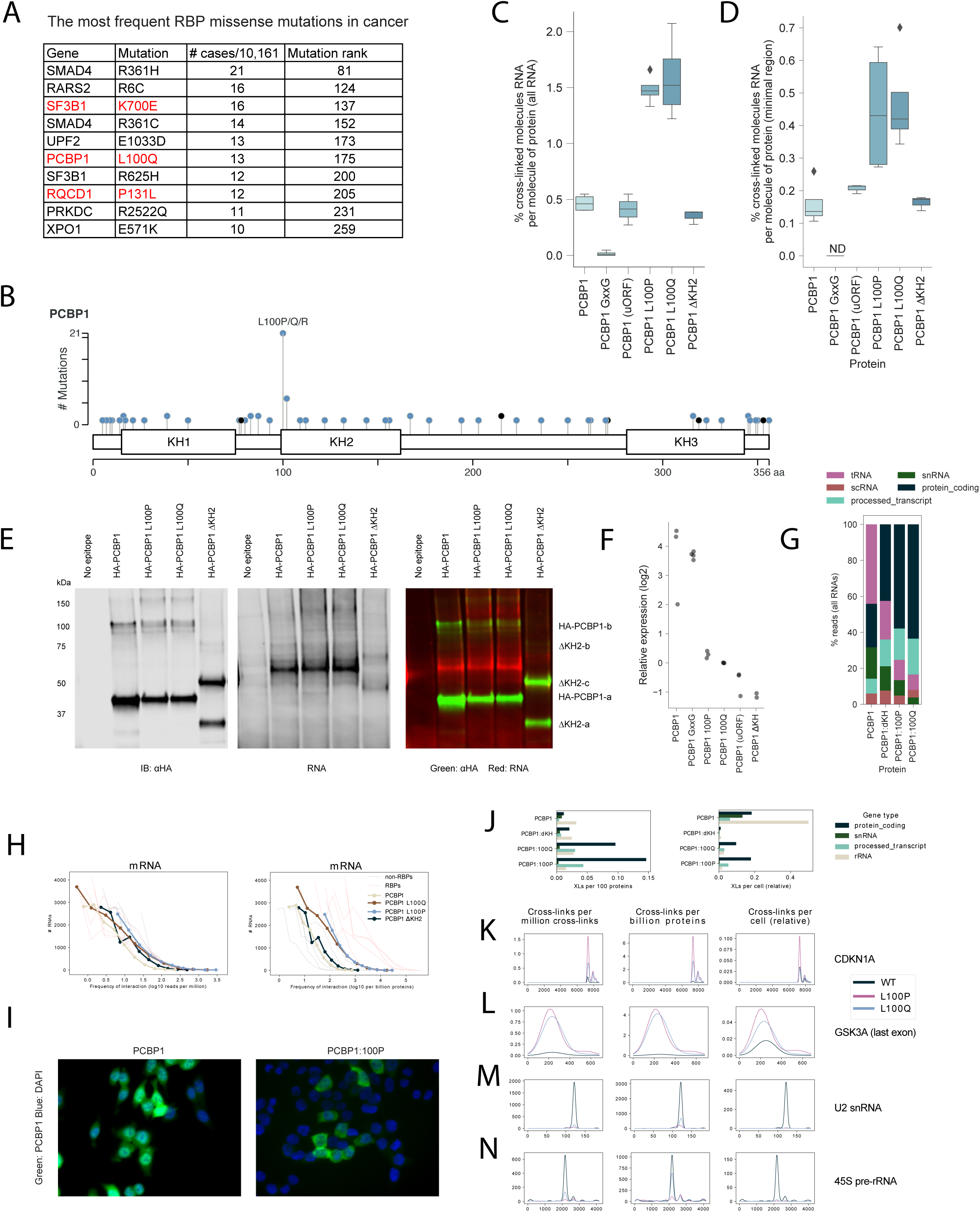
Cancer-associated missense mutations in RBPs investigated by easyCLIP. **A** The most frequent RBP missense mutations in cancer. Mutation rank is out of all mutations in TCGA data^15^. **B** Locations of missense (blue) and nonsense (black) mutations in PCBP1^15^. **C** Cross-link rates of wild-type and mutant HA-tagged PCBP1. **D** Cross-link rates of wild-type and mutant PCBP1, including only RNA signal in the minimal region. PCBP1 (uORF) represents wild-type PCBP1 expressed from a vector with a uORF to lower expression levels. **E** Visualization of immunopurified HA-PCBP1 by immunoblot, and cross-linked RNA by L5 adapter fluorescence. The right panel, viewing RNA and protein together, does not exactly match the RNA visualized alone because RNA degrades during the immunoblot process. RNA was always quantified immediately after transfer. **F** Expression levels of wild-type and mutant HA-PCBP1 in lysate, as determined by immunoblot and normalized to PCBP1 L100Q. **G** Distribution of reads between the indicated RNA types for PCBP1 and PCBP1 mutants. **H** Histograms of PCBP1 binding at mRNA on a *per read* or *per protein* basis. Non-RBPs are shown as gray lines, and RBPs are shown as red lines. **I** PCBP1 shifts from the nucleus to the cytoplasm in PCBP1 L100P mutants. **J-N** PCBP1 and L100P/Q PCBP1 mutant binding at the indicated RNAs. Signal was smoothed by fitting to a Gaussian kernel to enable visualization. GAPDH is shown in pre-mRNA form (the genomic locus). The left column is reads per million; the middle, cross-links per billion proteins; the right, cross-links per cell. The scale in the rightmost column is arbitrary and is not comparable between the different RNAs.

PCBP1 is both transcription factor and RBP, both nuclear and cytoplasmic, and highly multifunctional beyond RNA-binding^1^^6,17^. As a result, PCBP1 was expected to cross-link less than the average RBP. The cross-link rate of wild-type PCBP1 was indeed higher than non-RBPs, but lower than other RBPs (Figure 6C and D). To test if cross-linking was specific, GxxG loop mutations were introduced in all three KH domains of PCBP1, which remove the affinity of KH domains for RNA while allowing the domains to fold properly^18^. “GxxG PCBP1” no longer cross-linked to RNA (<0.01%, Figure 6C).

The effects of the PCBP1 L100 mutation were next examined. The first and second KH domains of the closely related protein PCBP2 form a pseudo-dimer, in which the β1 and α3 elements of both KH1 and KH2 bury hydrophobic residues against the other domain to form an intramolecular dimer^19^. L100, in β1 of KH2, is part of this dimerization surface^19^, suggesting L100 mutants might alter conformation to impair RNA-binding.

Surprisingly, the opposite effect was observed: L100P/Q PCBP1 was three-fold *more* cross-linked to RNA (Figure 6C-E). L100P/Q PCBP1 was dramatically destabilized (Figure 6F, Figure S11). Expressing PCBP1 from a vector containing an upstream ORF that lowered expression to below that of L100P/Q PCBP1 (Figure 6F) did not substantially increase cross-link rate (Figure 6C, D), ruling out expression levels as the cause of differential RNA-binding. These results indicate most of the wild-type protein is not bound to RNA in HCT116. Interestingly, if the entire KH domain containing L100 (KH2) is removed, cross-linking was approximately the same as wild-type (Figure 6C, D), yet ΔKH2 PCBP1 was also destabilized (Figure 6F, Figure S11).

L100P/Q mutants had a much smaller fraction of reads mapping to snRNA (Figure 6G), and on a per protein basis, L100P/Q greatly increased its association with mRNA (Figure 6H). It was therefore hypothesized that L100P/Q PCBP1 was more cytoplasmic than wild-type PCBP1, which was confirmed by microscopy (Figure S12, Figure 6I). ΔKH2’s location was unaltered (Figure S12).

The quantifications done by easyCLIP allow for new insight, as three different views of RNA-protein interactions are enabled (Figure 6J-N). Binding to snRNA by L100 PCBP1 is reduced per protein, but on a per cell basis it is clear the snRNA association of PCBP1 collapses in the L100P/Q mutants (Figure 6J, M). Although mutant PCBP1 interacts more often with mRNA per protein, per cell it is similar (Figure 6J, K, L). We note that the increase in GSK3A association is strong enough to overcome the effect of reduced abundance (Figure 6L). Altogether, Figure 5 and Figure 6J-N highlight the complexity of RNA-protein interactions, and how misled one might be if restricted only to analyzing CLIP data on the traditional basis of read distributions.

## Discussion

easyCLIP provides a general method for estimating RNA-per-protein cross-link rates. easyCLIP is easy, fast, reliable, and efficient. It provides direct visualization of the success of library preparation steps, allows multiplexing based on two adapters, and determines ligation efficiency. A major limitation to this approach is its reliance on UV cross-linking as a proxy for *in vivo* interactions^20^.

These PCBP1 results are consistent with a model where the L100P/Q mutations impair the stabilizing effect of KH2 and have a gain-of-function for KH2 with regards to location and RNA-binding. To the author’s knowledge, this is the first time a disease-associated mutation in an RBP has resulted in increased RNA-association.

PCBP1 protein is often down-regulated in cancer, which aids in tumorigenesis^21^. It is likely that the L100P/Q mutations contribute to tumorigenesis at least partly by destabilizing PCBP1. However, L100P/Q is only observed at high frequency in colon and rectal adenocarcinoma and down-regulation cannot explain the selection of a specific missense mutation. PCBP1 has been proposed to suppress tumors by binding mRNA and stabilizing tumor suppressor mRNAs, repressing translation of oncogenic mRNAs, and inhibiting oncogenic splicing^21^. The changes per cell we observe, however, indicate that while the landscape of PCBP1-RNA interactions is radically altered, the number of mRNA-PCBP1 complexes are similar with L100 mutants, and rather point to either changes in regulatory effect or a loss of function in splicing.

## Methods

The easyCLIP protocol is described in File S1 with additional information in Supplementary Methods. Full methods are in the Supplementary Methods section. High-throughput sequencing data is under the GEO accession GSE131210.

## Supporting information

Protocol

Description of datasets

Counts

Significance values

Peak locations

## Acknowledgments

We thank Brian Zarnegar for reagents, and input on experiments and their interpretation. We thank Amin Zia for performing analysis of TCGA data to identify missense mutations in RBPs in cancer. We thank Zurab Siprashvili and Yuning Wei for assistance. The SF3B1 sequence was obtained from a vector produced by Angelos Constantinou, provided by Marc-Henri Stern. Funding was provided by the NIAMS/NIH grant 1F32AR072504 to D.F.P., NIAMS/NIH grants AR45192, AR007422 and AR49737 to P.A.K., and by a USVA Merit Review grant to P.A.K. Some data was generated on an Illumina HiSeq 4000 purchased with funds from NIH award S10OD018220 by SFGF at Stanford.

## Author Contributions

D.F.P. wrote the software and conceived, designed, performed and analyzed all experiments. P.K. and D.F.P. planned experiments and wrote the paper.

## Supplementary material

### Supplementary files

**File 1** The full easyCLIP protocol and oligonucleotide sequences.

**File 2** Description of high-throughput sequencing datasets included in this study.

**File 3** Raw counts, counts per million reads, and counts per ten billion proteins for all proteins.

**File 4** P values for all proteins across all RNAs, determined by negative binomial fits to random non-RBPs in all cases.

**File 5** Peak locations for all proteins in all RNAs.

## Supplementary figures

**Figure S1.**
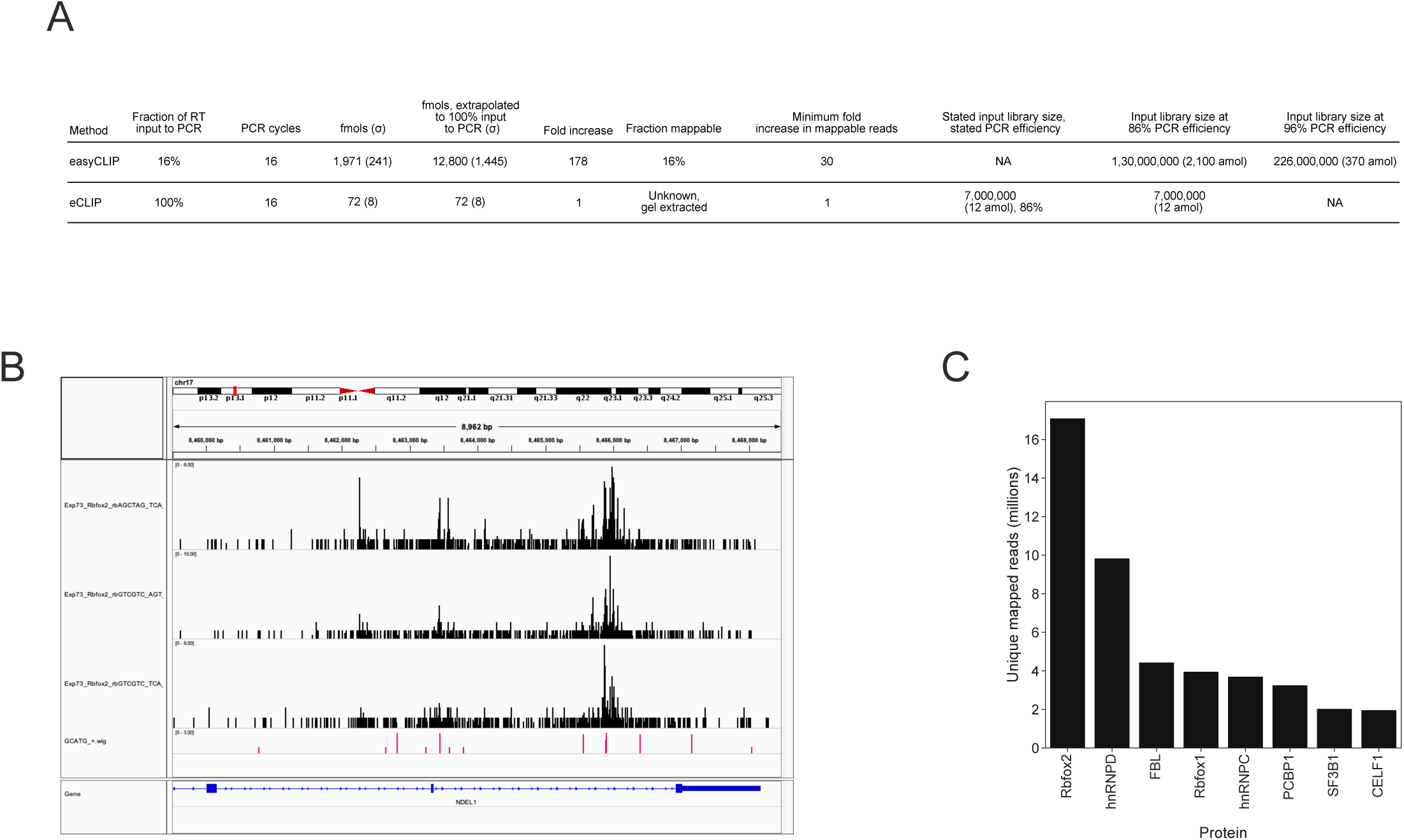
**A** Comparison of easyCLIP with eCLIP. The comparison used the same amount of the same anti-RBFOX2 antibody, the same cell line, and the same number of cells to perform easyCLIP on RBFOX2. eCLIP produced 72 fmols of library after 16 PCR cycles per replicate, as reported^1^, while easyCLIP produced ~13,000 fmols of library after the same number of cycles per replicate (n=3, extrapolating from PCR amplification of 16% of RT reactions). E.L. Van Nostrand *et al.* state that at 100% PCR efficiency their largest replicate would reach 100 fmol after 13 PCR cycles^1^. Dividing 100 fmol by 2^13^ gives an initial library size of 12 amol for eCLIP (7 million molecules) and a PCR efficiency of 86%. The subsequent information on RBFOX2 mapping in E.L. Van Nostrand *et al.*^1^ could not have come from this benchmark sample, as the authors report 85% unique reads at 20 million reads sequencing depth, impossible with a starting library of 7 million. eCLIP performed a size selection on their amplified library before sequencing, so the fraction of the input 12 amol that was usable is unknown. This easyCLIP sample did not undergo size selection before sequencing, resulting in many inserts too small to map, but 16% of reads were mappable. If easyCLIP PCR was 96% efficient (*vs* 86% for eCLIP), the starting pool would still be 370 amols. RBFOX2 data was obtained without substantial optimization (three RNAse concentrations were tried) – suggesting RBFOX2 does not represent an optimal case but a typical case. **B** Snapshot of the IGV browser viewing easyCLIP RBFOX2 reads at the same NDEL1 locus as shown in E.L. Van Nostrand *et al.*^1^ Figure 1D, showing identification of the same binding sites. Note that the scale bar in E.L. Van Nostrand *et al.* is reads per million, while the scale here is simply raw reads. Reads are placed according to their 5’ end location with a single nucleotide width. The GCATG_+.wig tract in red shows the location of GCATG motifs (the Rbfox2 binding site) on the plus strand, with a value of one placed on GCATG, a value of two placed on TGCATG (a preferred form of the motif), and allowing values to sum. **C** Unique mapped reads for eight RBPs. All data was obtained from 293T cells except PCBP1 was obtained from the colon cancer cell line HCT116. Cellular inputs ranged from below 10 million cells (hnRNP C, exact number not recorded), to 10 million (one RBFOX2 replicate), to 20 million (two RBFOX2 replicates), to a maximum of a 15 cm plate. RBFOX2, FBL, and hnRNP C libraries were obtained from antibodies to the endogenous proteins, the others were obtained from FLAG tag purifications from either constructs either integrated at the AAVS1 locus (PCBP1) or transiently over-expressed from a pLEX vector (the others).

**Figure S2.**
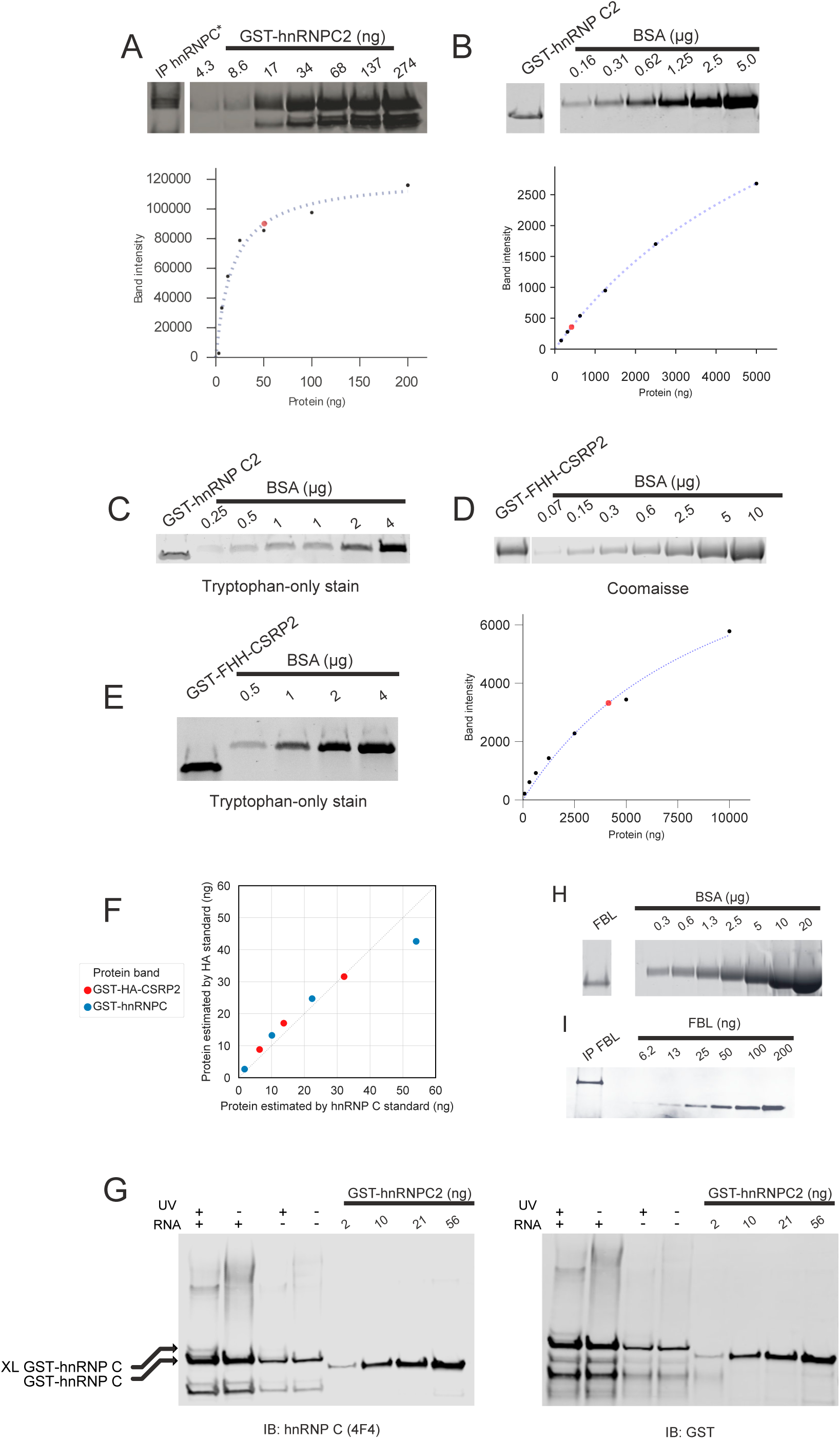
Quantification of purified recombinant protein and its application to absolute quantitation of immunopurified protein in CLIP. **A** Quantification of immunopurified endogenous hnRNP C using a GST-hnRNP C standard. The gel is a western blot probed with antibodies to hnRNP C. Endogenous hnRNP C is smaller than GST-hnRNP C but is shown at the same vertical position in this panel as GST-hnRNP C for visualization. In the graph, black dots represent GST-hnRNP C standards, the blue line is a best fit hyperbolic curve, and the red dot is immunopurified endogenous hnRNP C. **B** Quantification of purified GST-hnRNP C expressed in *E. coli*. GST-tagged hnRNP C was purified from *E. coli* using glutathione resin, and then run next to a standard curve of BSA protein on an SDS-PAGE gel. Gel was stained with Coomaisse and fluorescence measured at 700 nm. In the graph, black dots represent BSA standards, the dotted line is a fit hyperbolic curve, and the red dot represents the purified GST-hnRNP C, its position on the y-axis determined from the standard curve. The larger graph is focused on the lower quantities of GST-hnRNP C, while the larger graph is the same graph zoomed out to include all standards. **C** Quantification of GST-hnRNP C using a tryptophan-reactive dye (Bio-Rad Stain-Free Gel). Gel was subsequently stained with Coomaisse to determine Coomaisse staining of GST-hnRNP C and BSA was not biased. **D** Coomaisse quantification of purified, recombinant GST-FLAG-HA-His-CSRP2 (GST-FHH-CSRP2), the HA standard. CSRP2 was used in this construct because this fusion protein purifies in very high quantities. The hyperbolic curve fit is as in panel B. **E** Quantification of GST-FHH-CSRP2 using a tryptophan reactive-dye to test for a bias in Coomaisse-staining of the HA standard. No bias was observed. **F** Comparison of the quantification standards for HA and hnRNP C. Dilutions of each standard were run on the same gel and western blotted for GST. The standard curve of each protein stock was used to estimate the quantities of the other stock. The proximity of the dots to the 45° line indicate a good agreement. **G** The 4F4 anti-hnRNP C antibody shows little bias between cross-linked and non-cross-linked hnRNP C. Recombinant GST-hnRNP C (made in-house) was incubated with a poly(U)_10_RNA oligonucleotide (IDT) and UV cross-linked. The resulting mixture, along with GST-hnRNP C (Abnova) standards was run on a denaturing SDS-PAGE gel and transferred to a nitrocellulose membrane for immunoblotting against hnRNP C (4F4) or GST. No significant difference between anti-GST and anti-hnRNP C antibodies in the ratio of cross-linked to non-cross-linked hnRNP C was observed. **H** Coomaisse quantification of purified, recombinant FBL. Purified FBL protein (Prospec, enz-566) was comprised of FBL amino acids 83-321 with an added 23 amino acid tag added, and the FBL antibody (Bethyl, A303-891A) was made against an immunogen between amino acids 271-321 of FBL. As a result, the purified FBL runs faster than endogenous FBL, but both share the entire immunogen used for immunoblotting. **I** Immunoblot quantification of immunopurified FBL using the recombinant FBL visualized in panel H.

**Figure S3.**
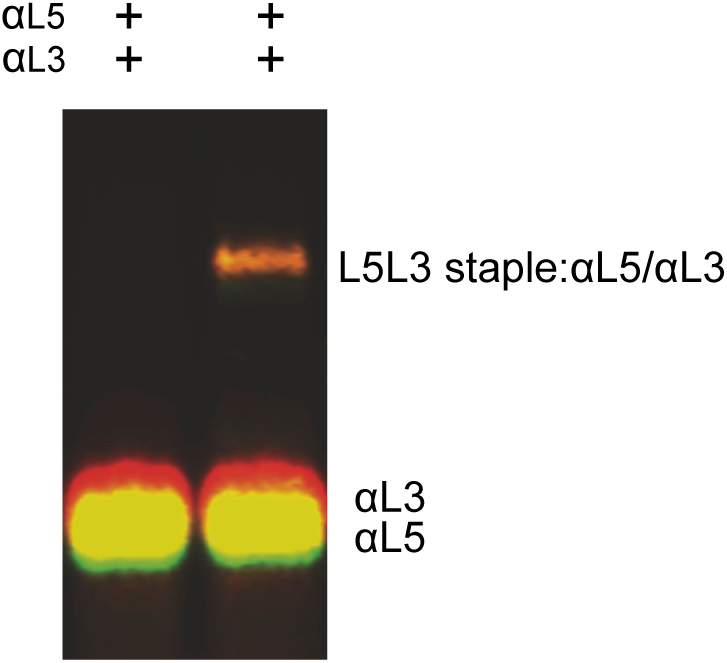
A staple oligonucleotide may be used to shift the antisense oligonucleotides in Figure 2D in a single molecule to determine relative fluorescence and control both adapter quantifications to a single complex.

**Figure S4.**
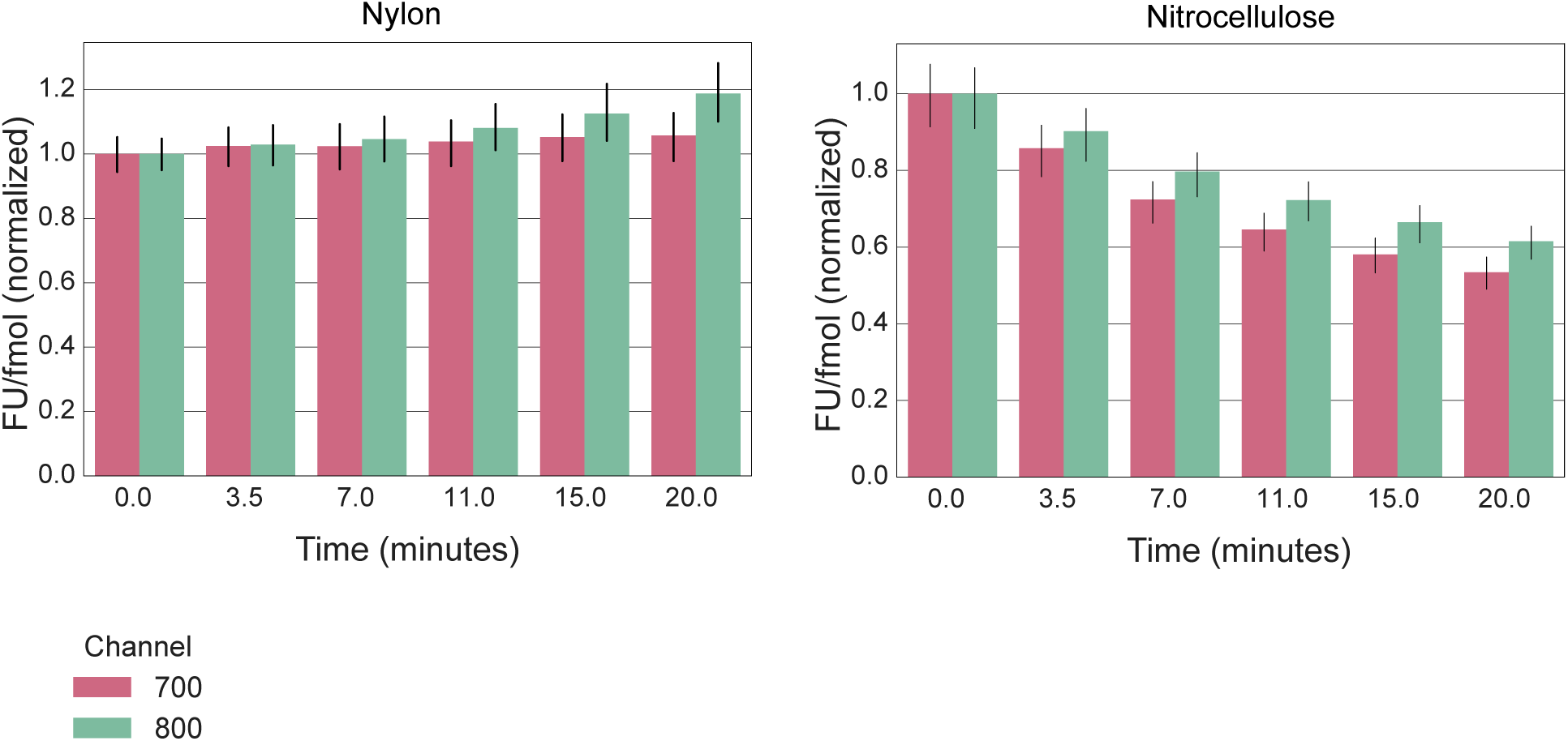
Fluorescence on nylon and nitrocellulose for dot blots of αL3 and αL5 labelled respectively with IR680RD and IR800CW. Signal remains high on nylon, but decays on nitrocellulose

**Figure S5.**
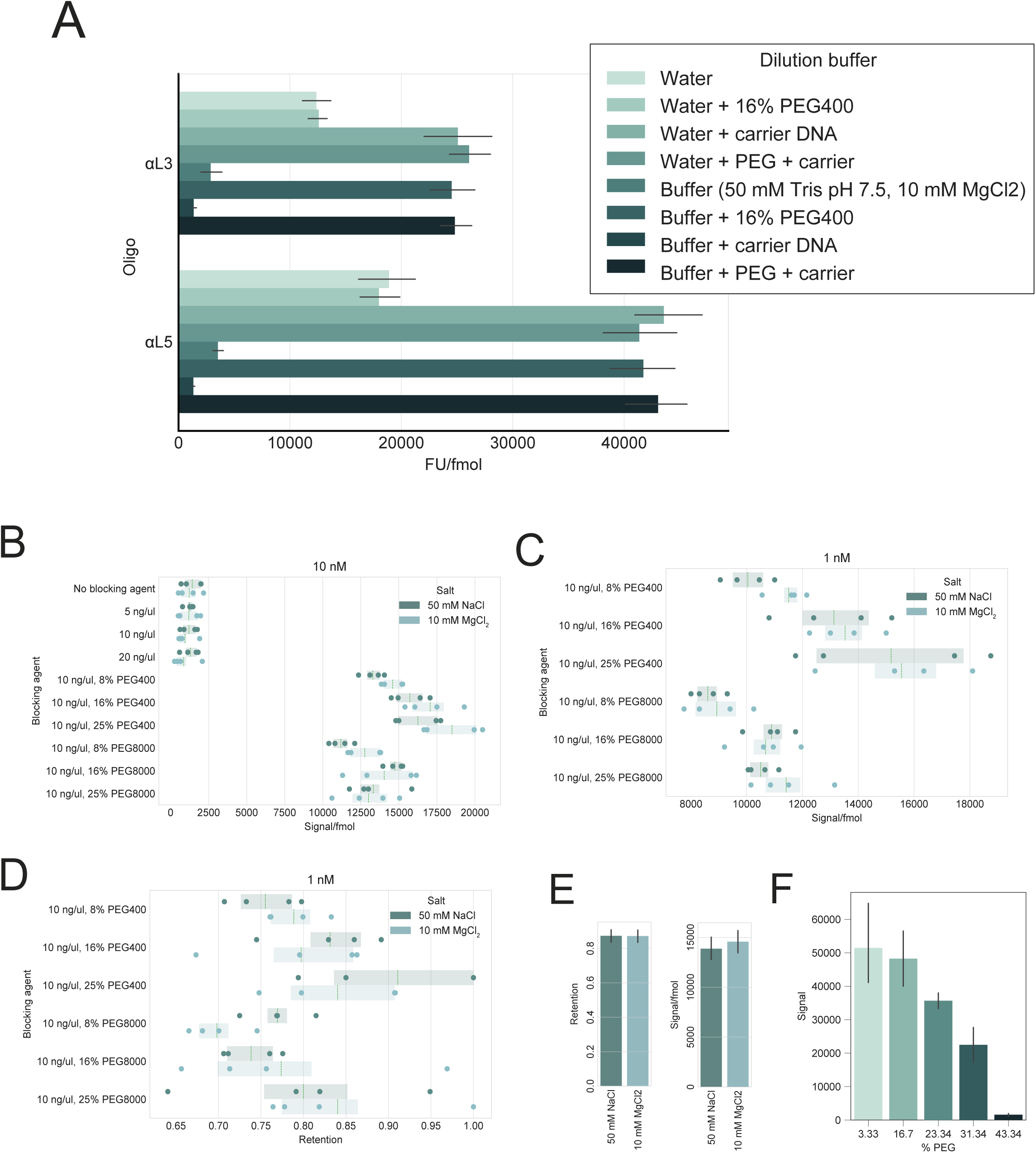
Developing a method to quantify low fmol amounts of adapter. **A** The choice of dilution solution has a large effect on fluorescence. An equimolar mixture of αL3 and αL5 was dilute to 1 nM in the indicated solutions. 2 µL (2 fmols) of diluted oligonucleotide were then dot blotted on nylon and fluorescence measured on a Li-Cor scanner. Carrier DNA was an equimolar solution of 10, 15, and 35 nucleotide poly(A) oligonucleotides. **B** Fluorescence per fmol of αL3 oligonucleotide after diluting to 10 nM in 50 mM Tris pH 7.5 with the indicated salts and blocking agents. Carrier DNA was an equimolar solution of 10, 15, and 35 nucleotide poly(A) oligonucleotides at the indicated ng/µL concentrations. All PEG solutions had 10 ng/µL carrier DNA. Carrier DNA is not sufficient to block signal loss upon dilution. Both monovalent and divalent salts had similar effects. PEG400 and PEG8000 both preserved signal, and higher concentrations generally worked better. **C** The 10 nM solution in panel B was diluted to 1 nM. PEG400 leads to slightly higher fluorescence than PEG8000. Solutions lacking PEG are not depicted due to low signal to noise ratios. **D** Retention of signal during a 10-fold dilution. Retention is the fluorescence per fmol of the 1 nM solution divided by the fluorescence per fmol of the 10 nM. The choice of salt has no consistent effect. Higher PEG concentrations are better blocking agents. PEG400 and PEG8000 have a similar performance as blocking agents. **E** The choice of 50 mM NaCl or 10 mM MgCl_2_ has no effect on oligonucleotide loss during dilution (retention) or on signal per fmol. **F** It is safe to run DNA duplexes on 20% polyacrylamide TBE gels (NuPAGE, 12 well, ThermoFisher) at 16.7% PEG400, but higher concentrations lead to fluorescence loss in the duplex, probably due to unfolding of the DNA duplex.

**Figure S6.**
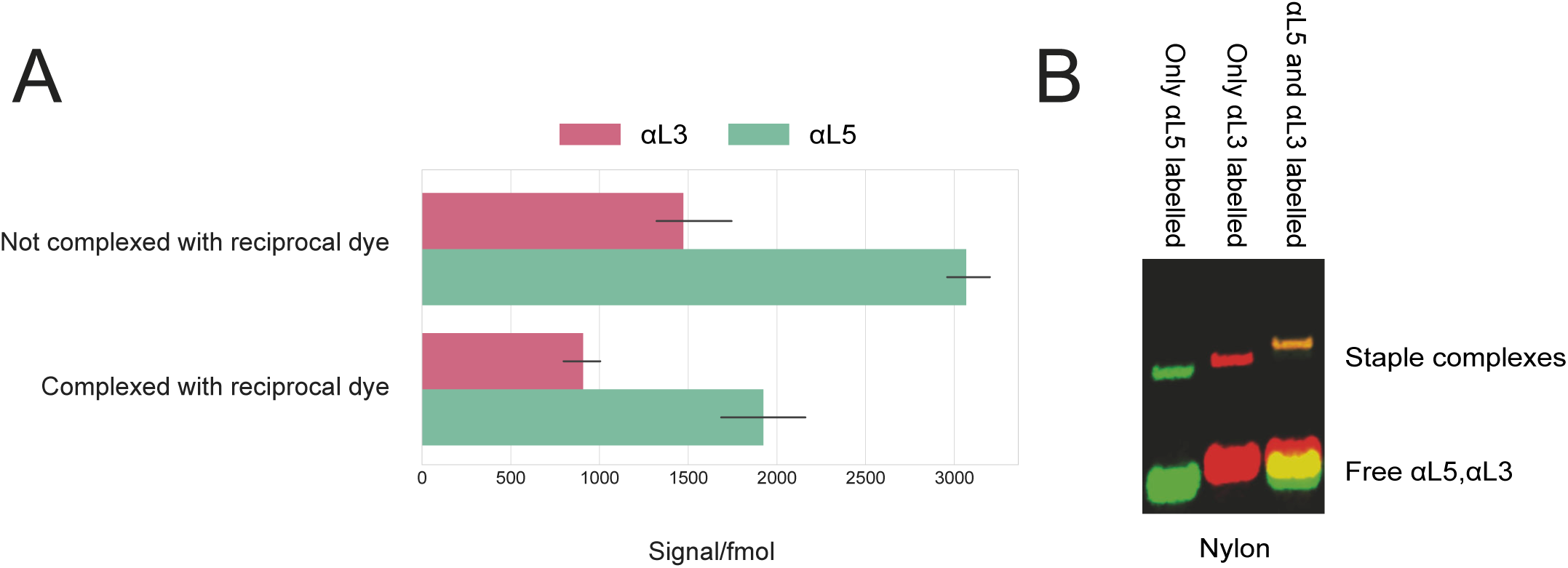
Signal interference between IR800CW and IR680RD dyes. **A** The IR800CW and IR680RD dyes decrease in fluorescence when tethered to the same complex. An excess of αL5 and αL3 were mixed with 50 fmol of an oligonucleotide bearing one copy each of the L5 and L3 sequences, termed the staple oligonucleotide. αL5 was paired with either labelled or unlabeled αL3 to determine the effect of tethering αL3 near αL5, and the reciprocal case was applied to αL3. Complexes were run on a TBE gel in TBEN buffer (0.5X TBE plus 50 mM NaCl) and transferred to a nylon membrane for quantification. **B** Labelled complexes always traveled higher on the gel (right panel). Each dye shifts ~6 nucleotides higher on a TBE gel.

**Figure S7.**
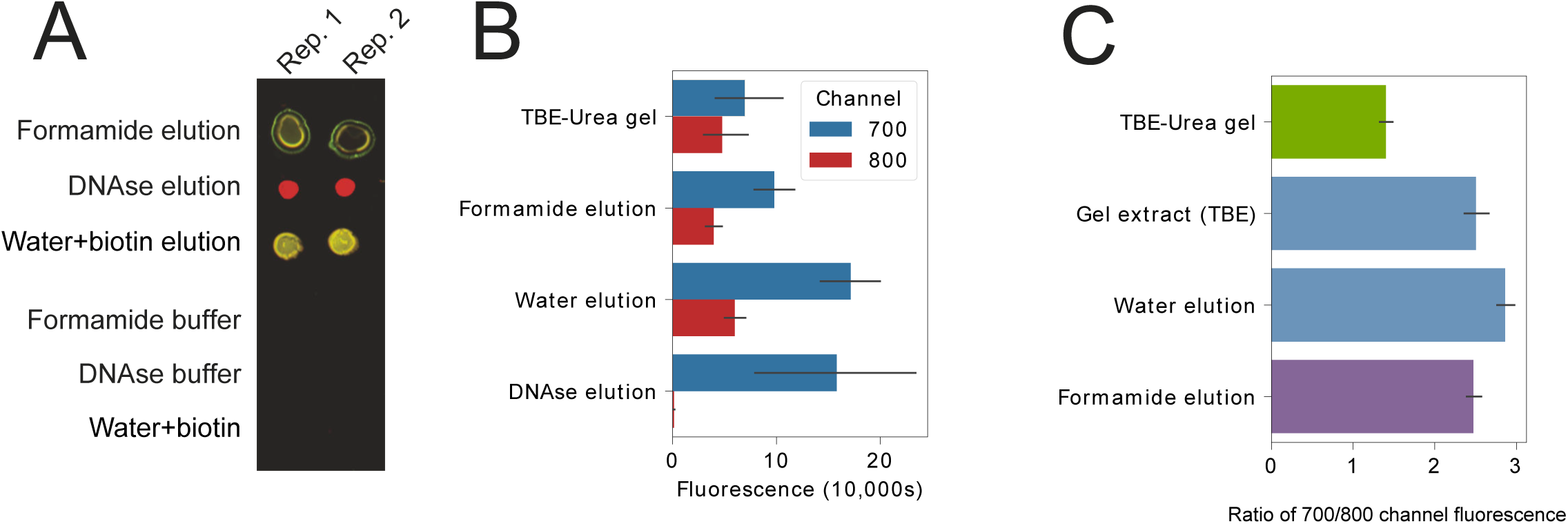
Performance of streptavidin elution methods. **A** L5 and L3 adapters were ligated together *in* vitro, run on a TBE-urea gel, gel extracted, purified using streptavidin beads (MyOne C1, ThermoFisher), and then eluted by the indicated method. This image shows an example of eluates dot blotted on nitrocellulose. Note the peculiar shape of formamide dots. No fluorescence is observed in buffer alone. Water+biotin elution used 100 nM biotin. Formamide elution was 95% formamide with 10 mM EDTA (as suggested by ThermoFisher, who state elution is >95% by this method). DNAse elution used an excess of DNAse I (Ambion) in the buffer supplied by the manufacturer. **B** Fluorescence quantification of the same linker-linker dimers depicted in panel A after each elution method. “TBE-urea gel” indicates fluorescence in the TBE-urea gel before extraction and streptavidin purification. Heating in water with 100 µM biotin was effectively complete, as it yielded similar L5 (700 nm) fluorescence as DNAse elution, which is likely to be complete, and similar fluorescence overall as formamide elution, which is complete according to the manufacturer (ThermoFisher). **C** Water, formamide and TBE-urea gels all affect relative L5/L3 fluorescence (IR680RD/IR800CW). The ratio of dye molecules is 1:1 in all cases, as all cases represent linker-linker dimers.

**Figure S8.**
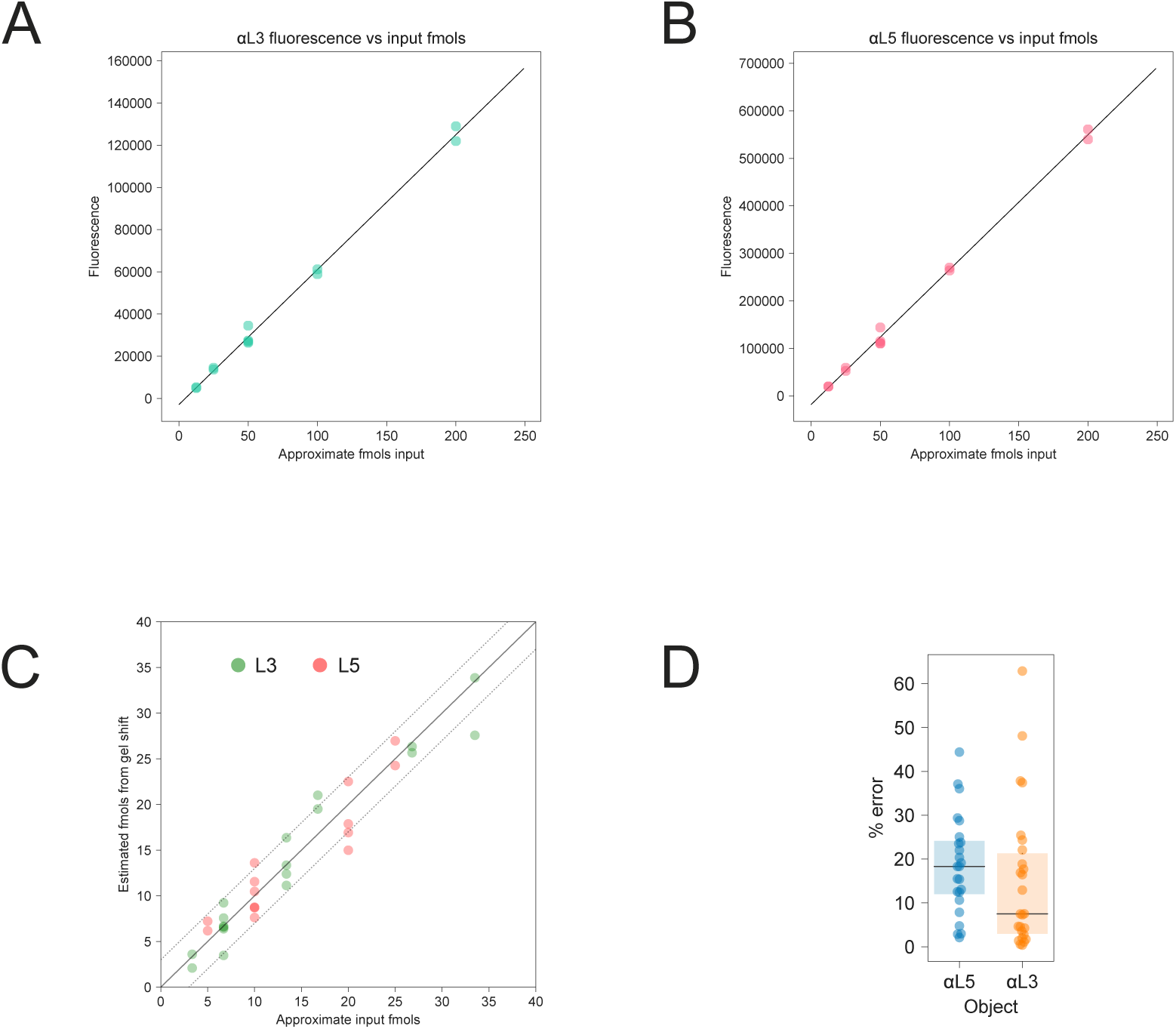
Model-fitting and testing of an anti-sense oligonucleotide shift method of adapter concentration. **A** Fluorescence of the αL5 oligonucleotide in the staple-αL5-αL3 complex as a function of staple oligonucleotide quantity. Signal fits to a linear model (solid line). **B** Fluorescence of the αL3 oligonucleotide in the same complexes as A. Signal is again highly linear (solid line is a linear fit). **C** Known concentrations of L5 and L3 adapters and staple oligonucleotide were shifted by αL5 and αL3 and a fit to a linear model. As with staple oligonucleotides, data is linear: the solid line represents a perfect fit, dashed lines represent + or – 3 fmols. **D** Error in the estimates made in panel C. The method is reasonably accurate, with average errors around 20%. The parameters (slope and intercept) from panel C were then used to estimate oligonucleotide concentrations for ligation efficiency determinations, after applying a scaling factor based on the fluorescence of αL5/ αL3 oligonucleotides in 50 fmol staple complexes. The calculation is described in github.com/dfporter/easyCLIP/doc/ in the README_fluorescence.md file.

**Figure S9.**
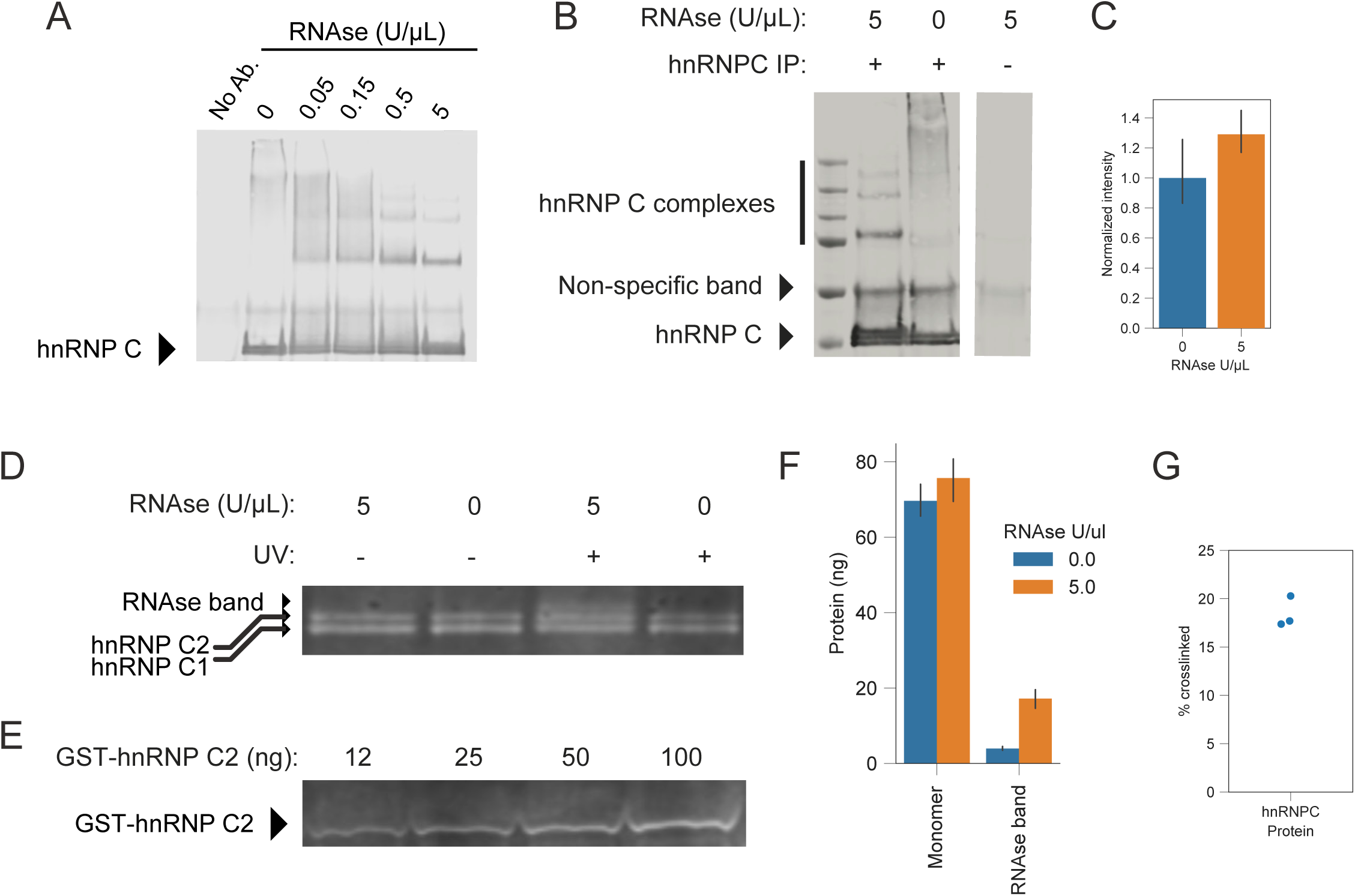
Quantification of cross-link rates for endogenous hnRNP C by immunoblot shift. Cells were UV cross-linked cells then hnRNP C was immunopurified. The change in western blot signal corresponding to monomeric hnRNP C was compared between RNAse concentrations (panels A-C). Because this change in signal is specifically for what can be collapsed with RNAse to monomeric hnRNP C, not for the un-collapsible higher molecular weight complexes spread throughout the lane, it should agree with the cross-linking number derived from dividing the RNA quantified in the minimal region by the monomeric hnRNP C signal (Figure 4C) and be lower than that derived from all RNA across the gel. **A** RNAse digestion series of immunopurified hnRNP C (immunoblot, anti-hnRNP C). **B** Example replicate of +/− RNAse gels used to quantify the amount of shifted hnRNP C. **C** Quantification of the amount of shifted immunoblot signal comparing +/− RNAse gel lanes, as in panel B. The change in western blot signal was ~20%, close to the 22% cross-link number from Figure 4C. A more exact comparison was then performed, deriving the amount of hnRNP C protein dependent on both UV cross-linking and RNAse-digestion by absolute quantification of a western blot (panels D-F). **D** Gel used for absolute quantification of UV- and RNAse-depending monomeric hnRNP C signal. **E** Standards used for absolute quantification of gel data as in panel D. **F** Quantification of the absolute amount of protein present in the bands in replicates like that in panel D. **G** The amount of hnRNP C cross-linked to RNA that is collapsible into the monomeric hnRNP C band, as determined by the absolute quantification data in panel F. This method also gave a cross-link rate of ~20%, again similar to the 22% observed in Figure 4C. It was concluded that this method of determining cross-link rates using absolute quantification of RNA and protein (Figures 2 and 3) was reasonably accurate. This verification was only possible for hnRNP C because of its very high cross-link rate and small size.

**Figure S10.**
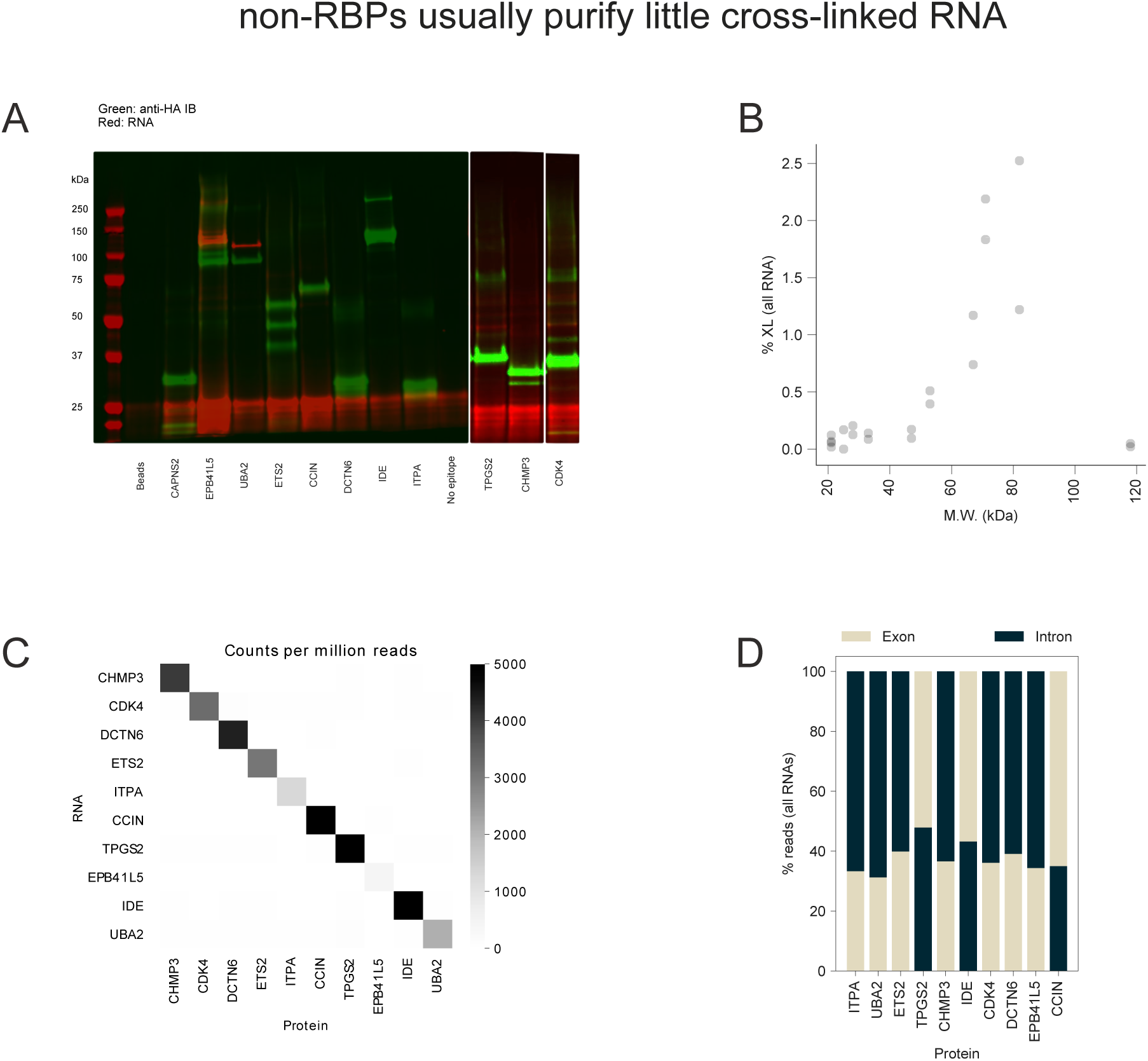
**A** Purification of randomly selected HA-tagged non-RBPs. Red represents L5 adapter fluorescence, and green anti-HA immunoblotting. **B** Total purified cross-linked RNA positively correlates with protein size for randomly selected non-RBPs. **C** Immunoblot and RNA visualization of the two non-RBPs that purified the most cross-linked RNA, UBA2 and EPB41L5, shows cross-linked bands running a little higher than the minimal region. **D** Read counts (per million reads) of the non-RBPs *vs* their own RNAs shows each non-RBP enriches for its respective RNA, a consequence of each non-RBP being expressed from a plasmid. This shows each library was generated from cells over-expressing the respective protein-of-interest, despite the fact that barcodes for multiple over-expression experiments were combined after each ligation. It also shows that if you express an RNA highly, it will show up in CLIP data, regardless of the purified protein. Counts were capped at 5,000 reads-per-million for visualization. Libraries for CAPNS6 were extremely small and were not included. **E** Distribution of reads between introns and exons in mRNA for randomly selected non-RBPs.

**Figure S11.**
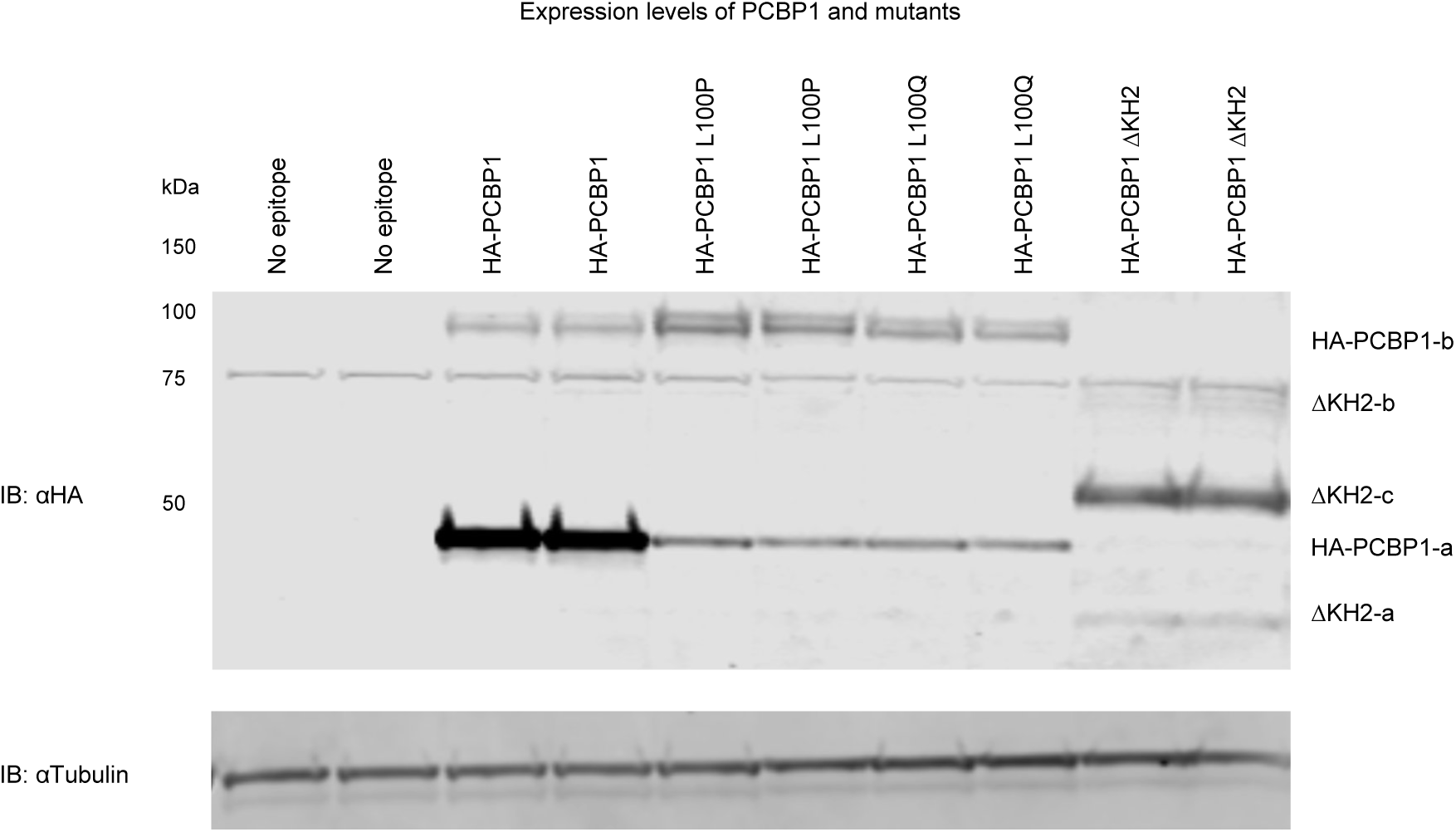
Expression levels of FH-PCBP1 and mutants in HCT116 cell lysate. The nature of the additional, higher molecular weight bands (b, c) is unknown.

**Figure S12.**
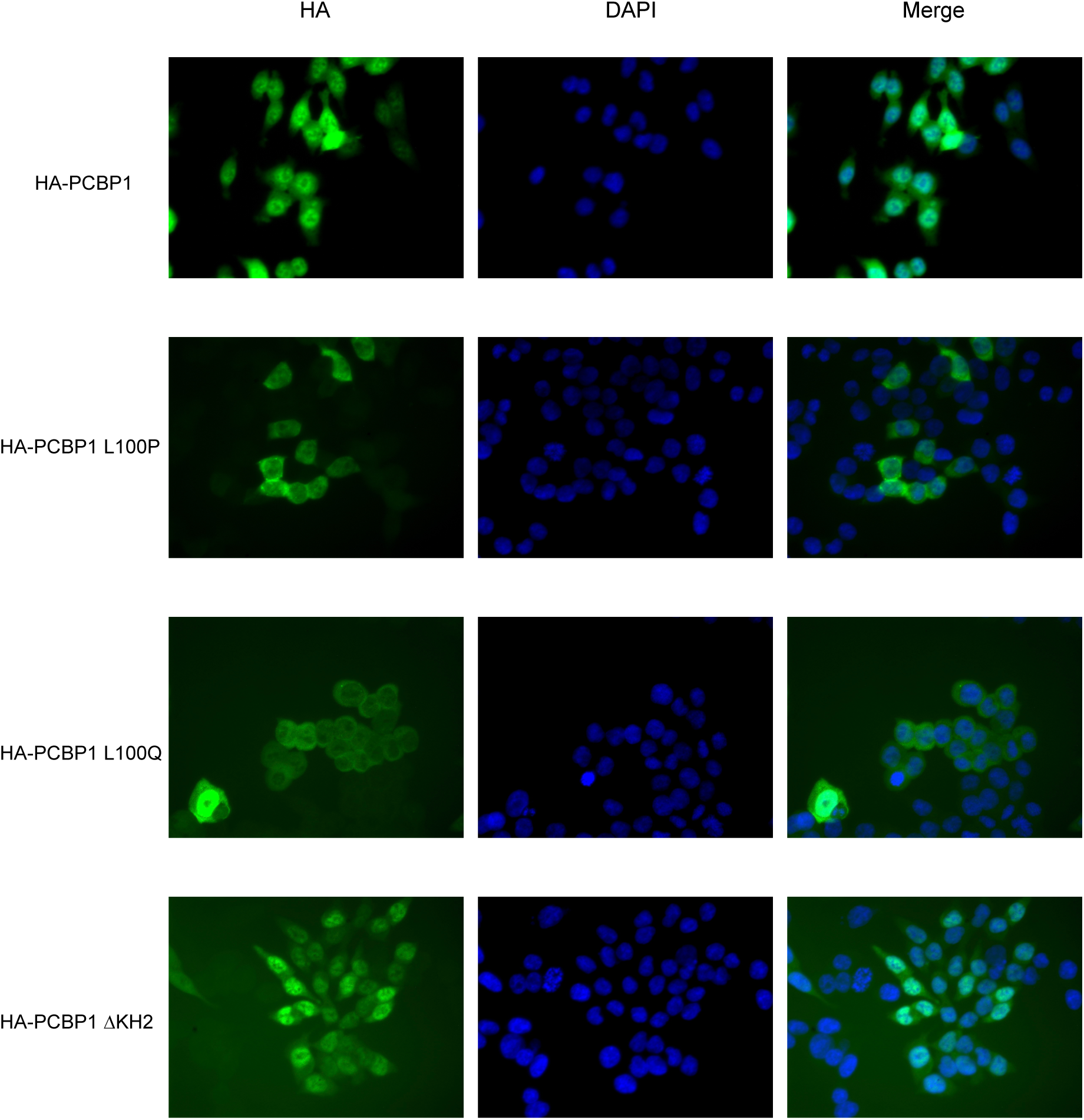
Microscopy of wild-type and mutant FHH-PCBP1 in HCT116 cells showing that L100P/Q mutants are less nuclear than wild-type or ΔKH2 PCBP1. All images were taken with the same settings (exposure time, *ect.*), on the same slide and day.

## Supplementary Methods

### L5 linker labelling

0.5 mg IRDye 680RD DBCO (LI-COR, 429 nmol) was resuspended in 42.9 µL PBS for a concentration of 10 mM. The L5 linkers (Azide-DNA-RNA oligonucleotides) were ordered from IDT and resuspended in PBS. Oligonucleotides were run through a Zymo RNA-clean-and-concentrator kit (purification was required for labelling), using ~14 µg oligonucleotide per column and eluting at ~1 mg/mL (~85 µM) in water. 5 µL of 10 mM dye (~50 nmol) was added to 10-150 µg purified oligonucleotide (~1-12 nmol) in PBS for a total volume of 200 µL and reacted for 2 hours at 37°. Oligonucleotides were then run again through a Zymo clean-up kit and eluted in water. During column purifications, washes were performed using an 85% ethanol in water solution made fresh each time, in place of the kit’s wash buffer. Concentrations were determined by A260 ratio using an approximate ε=368,050 M^−1^. Oligonucleotides were diluted to 10 nM in ligation buffer (50 mM Tris pH 7.5, 10 mM MgCl_2_, 16.7% PEG400), 1 µL was blotted onto a nylon membrane, and fluorescence was measured in an Odyssey CLx machine (LI-COR). This was typically ~15,000 fluorescence units per fmol for full labelling.

### AAVS1 microscopy of PCBP1 integrants

4-well plastic chamber slides (Lab-Tek Permanox, Sigma #C6932-1PAK) were coated with 0.01% poly-L-lysine (Sigma #P4707) for 15 minutes, then washed twice with PBS, left dry for 5-30 minutes, and then either stored under PBS or used immediately. HCT116 cells were plated at <20% confluency and grown at least 24 hours before staining. Cells were washed 1-2 times with PBS, then fixed for 10 minutes in 4% formaldehyde (in PBS) at room temperature, rinsed three times with PBS, and then permeabilized with PBS containing 0.5% Triton X-100 and 10% goat serum. After permeabilization, cells were stained for 1 hour at room temperature with the primary antibody at 1:200 dilution in PBS containing 0.05% Triton X-100 and 1% goat serum. After staining, cells were washed three times with PBS containing 0.05% Triton X-100, then 2-3 times in PBS without detergent, and the slide chamber removed. After letting the cells dry for a few minutes, one drop of DAPI mounting solution was added to each well and a coverslip was added and sealed with acetone.

### AAVS1 integration

~2 µg repair template and ~1 µg Cas9/guide RNA plasmid were transfected using lipofectamine into 6-well plates containing ~300,000 cells each. Two days later, puromycin was added to 1 µg/mL and selection continued for at least 10 total days. To determine expression levels, 10 µg to 80 µg of clarified lysate in 1-8 µL of CLIP lysis buffer (typically 4 µL) was combined with 16 µL 1.6X LB (NuPAGE) and run on an SDS-PAGE gel. hnRNP C was immunoblotted using labelled anti-hnRNP C antibody (Santa Cruz, 798-conjugated) at 3 µL in 5-7 mL PBS blocking buffer (Licor), incubating for 30 minutes and washing with PBS for 20 minutes. To immunoblot for the HA tag, ~3 µL Rabbit anti-HA (COVANCE) in 5-7 mL blocking buffer, followed by ~3 µL IR680 or IR800 labeled Goat anti-Rabbit (Licor) in 5-7 ml were used.

### AAVS1 integrated FHH-tagged protein purification

15 µL anti-HA magnetic beads and 2-4 mg clarified lysate were used per immunopurification. Immunopurifications were carried out at 4° for 1 hour in 1 mL of CLIP lysis buffer.

### GST-tagged protein constructs

pGEX-6P-1 vector was digested with BamHI and CSRP2-FLAG-HA was cloned in using In-Fusion (Takara). Amplification primers for CSRP2-FLAG-HA were:

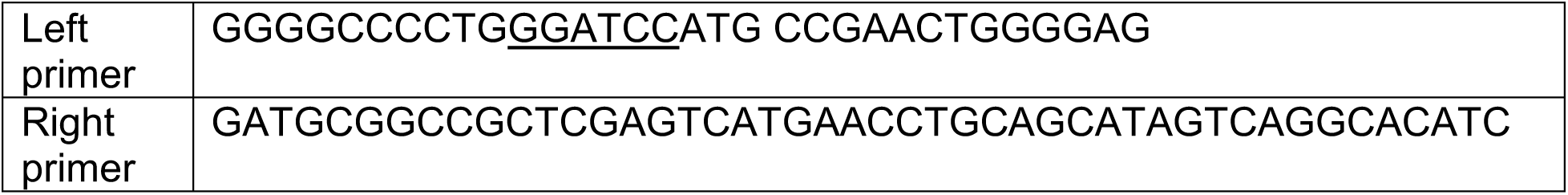

The GST moiety (and protease site) is 231 amino acids (26.8 kDa), and CSRP2-FLAG-HA is 217 amino acids (23.2 kDa), for a 448 amino acid (50 kDa) construct. This resulting sequence is given below, with CSRP2-FLAG-HA underlined (* denotes stop):

MSPILGYWKIKGLVQPTRLLLEYLEEKYEEHLYERDEGDKWRNKKFELGLEFPNLPYYI DGDVKLTQSMAIIRYIADKHNMLGGCPKERAEISMLEGAVLDIRYGVSRIAYSKDFETLKVDFLSKLPEMLKMFEDRLCHKTYLNGDHVTHPDFMLYDALDVVLYMDPMCLDAFPKLVCFKKRIEAIPQIDKYLKSSKYIAWPLQGWQATFGGGDHPPKSDLEVLFQGPLGS<u>MPNWGGGKKCGVCQKTVYFAEEVQCEGNSFHKSCFLCMVCKKNLDSTTVAVHGEEIYCKSCYGKKYGPKGYGYGQGAGTLSTDKGESLGIKHEEAPGHRPTTNPNASKFAQKIGGSERCPRCSQAVYAAEKVIGAGKSWHKACFRCAKCGKGLESTTLADKDGEIYCKGCYAKNFGPKGFGFGQGAGALVHSELEDYKDDDDKAGYPYDVPDYAAGS</u>*

The GST-hnRNP C construct (54 kDa) was cloned into the same site but did not include HA or FLAG tags. The resulting sequence is below:

MSPILGYWKIKGLVQPTRLLLEYLEEKYEEHLYERDEGDKWRNKKFELGLEFPNLPYYIDGDVKLTQSMAIIRYIADKHNMLGGCPKERAEISMLEGAVLDIRYGVSRIAYSKDFETLKVDFLSKLPEMLKMFEDRLCHKTYLNGDHVTHPDFMLYDALDVVLYMDPMCLDAFPKLVCFKKRIEAIPQIDKYLKSSKYIAWPLQGWQATFGGGDHPPKSDLEVLFQGPLGS<u>MASNVTNKTDPRSMNSRVFIGNLNTLVVKKSDVEAIFSKYGKIVGCSVHKGFAFVQYVNERNARAAVAGEDGRMIAGQVLDINLAAEPKVNRGKAGVKRSAAEMYGSVTEHPSPSPLLSSSFDLDYDFQRDYYDRMYSYPARVPPPPPIARAVVPSKRQRVSGNTSRRGKSGFNSKSGQRGSSKSGKLKGDDLQAIKKELTQIKQKVDSLLENLEKIEKEQSKQAVEMKNDKSEEEQSSSSVKKDETNVKMESEGGADDSAEEGDLLDDDDNEDRGDDQLELIKDDEKEAEEGEDDRDSANGEDDS*</u>

### GST-tagged protein purification

*E. coli* BL21 cultures transformed with pGEX-6P-1 were grown in 500 mL at 37° until OD600 ~0.8, at which time Isopropyl-1-thio-β-D-galactopyranoside (IPTG) was added to a final concentration of 0.5 mM, and cultures were grown for another ~1.5 h before harvesting. Cells were harvested by the method of S. Harper *et al.*^2^, namely centrifuging at 4,000 rcf for 20 min at 4°, resuspending in ~50 mL LB, and centrifuging again at 4,000 rcf for 20 min at 4°. Cell pellets were frozen in dry ice until purification. When thawed, the cell pellet was resuspended in 20 mL of lysis buffer (50 mM Tris, 10 mM β– mercaptoethanol, 50 mM NaCl, 5 mM EDTA, 1% Triton X-100, Roche protease inhibitor, 5% glycerol). Lysozyme was added very approximately to ~1 mg/ml, froze the pellet again in dry ice, thawed in a water bath, and lyzed by sonication. The lysate was clarified by centrifugation at ~21,000 rcf, 4°, for 15 min. 4 mL of 50% glutathione-agarose (Pierce) was washed with resin wash buffer (Dulbecco PBS with 10 mM β–mercaptoethanol), and then incubated at 4° in a 50 mL Falcon tube with clarified lysate for ~30 min before loading on a column. The column was washed with 50 mL of 4° wash buffer (Dulbecco PBS with 10 mM β–mercaptoethanol, 5% glycerol and Roche protease inhibitor). Samples were eluted in batch with three incubations at 4° with 1.5-2 mL elution buffer (100 mM Tris pH 8.0, 150 mM NaCl, 10 mM β–mercaptoethanol, 5% glycerol, 10 mM glutathione).

### GST-tagged protein quantification

Following the method of K. Janes^3^, BSA standards were run on a gel at 10, 5, 2.5, 1.3, 0.6, 0.3, and 0.15 µg, along with purified protein. Following the method of S. Luo *et al.*^4^, gels were washed for 10 minutes in water, stained for 10 minutes with staining buffer (50% methanol, 10% acetic acid, 0.02% Coomaisse R250) at room temperature, followed by destaining for 10 minutes with destaining buffer (40% methanol, 7% acetic acid), and washing twice for 10 minutes with water. A third wash was performed overnight. Protein was then visualized by scanning the 700 nm channel on a Licor Odyssey scanner. A hyperbolic curve of band fluorescence *vs* input protein weight was fit to BSA standards. Specifically, the parameters ‘a’ and ‘b’ in the equation y = a*x/(b+x), where ‘x’ is protein weight and ‘y’ is fluorescence, were fit using least-squares regression. This curve was used to determine the concentration of purified protein.

### Western blot protein quantification

Following the method of K. Janes^3^, purified GST-tagged protein standards were run alongside the samples to be quantified. Purified GST-hnRNPC2 and purchased FBL (Prospec, cat. enz-566) were diluted in protein dilution buffer (0.5X PBS, 0-5% glycerol, 0.05% Tween-20, 0.2 mg/mL BSA) to 20 ng/µL. Two-fold dilutions down from 20-100 ng/µL were made for a total of 8 concentrations; this solution was then delivered as 14 µL aliquots to multiple striptube aliquots and frozen at −80°. When running gels, 10 µL from each concentration were combined with 10 µL loading buffer (3.6X NuPAGE loading buffer with 10% β-mercaptoethanol), heated at 75° for 15 minutes, and loaded on a 4-12% NuPAGE gel. Standards were therefore present at ~1000-3 ng per lane. Immunoblotting against the HA epitope was performed with 1:3000 αHA conjugated to Alexa Fluor 488 and incubating for 1 hour at room temperature in PBS blocking buffer (LI-COR); images were taken in a GE Typhoon scanner (532 nm laser, 526SP filter, 500 PMT, 200 µm resolution). When small aliquots of immunopurification beads were loaded on a gel, BSA was first added to 0.2 mg/mL to prevent absorption.

### BCA

For BSA standards, 105 µL PBS was combined with 20 µL BSA (2 mg/mL stock) and 3 µL lysis buffer for the highest concentration of BSA, and 115 µL PBS, 10 µL BSA, and 3 µL lysis buffer for the second highest concentration. For lysate samples, 3 µL lysate was combined with 125 µL PBS. For both standards and samples, serial dilutions were made by a factor of three into PBS with 0.024% lysis buffer. Duplicate wells were used for each sample. 25 µL of each well was transferred to a second 96-well plate and combined with 200 µL working reagent (Pierce BCA kit, 50:1 A:B). Plate was incubated for 20-30 minutes at 37°. Absorbance was measured at 562 nm.

### Creation of cross-linked hnRNP C standard

Four replicates of 906-1600 µg of HCT116 lysate from cross-linked cells was added to ~20 µL Protein G Dynabeads (ThermoFisher Cat #10003D) coupled with 25 µL (5 µg) anti-hnRNP C (4F4) antibody per replicate. Immunoprecipitation was carried out at 4° for ~1 hour, followed by the standard easyCLIP protocol for cross-link rate determination. The RNAse digestion was performed with half of the samples treated with 0.1 U/µL RNAse ONE for 10 minutes, and the other half of the samples treated with 0.05 U/µL RNAse ONE for ~5 minutes. The PNK reaction was 14 minutes at 37°. The ligation was performed overnight (17 hours) with 20 pmol L5 (barcode 23), and 2 µL high concentration T4 RNA ligase (NEB). Samples were combined, and ~20 aliquots comprising 2.5% of the beads (~10 ng hnRNP C each, ~400 ng total purified) in ~15 µL 1.6X NuPAGE buffer were frozen in dry ice and kept long term at −80°. Immunoblotting was performed with ~1:3000 αhnRNP C conjugated to AF790 (Santa Cruz Biotechnology, sc-32308 AF790), which is visible on the 800 nm channel in a LI-COR Odyssey scanner, in PBS blocking buffer (LI-COR) for ~1 hour at room temperature.

### Sequencing library creation: hnRNP C and FBL

HEK293T cells were grown to 30-90% confluency in petri dishes in DMEM with 10% Fetal Bovine Serum, media was removed by vacuum, cells were washed with 4° PBS, and UV cross-linked (254 nm) in 10 cm or 15 cm plates in a Stratalinker at 0.3 J/cm^2^. After cross-linking, 1 mL 4° lysis buffer (15 cm plates) or 0.5 mL lysis buffer (10 cm plates) was added to each plate, cells were harvested with a rubber spatula and frozen in dry ice. CLIP lysis buffer was as in Zarnegar *et al.*^5^, except the concentration of Triton X-100 was 1% (see File S1 for all buffers used for CLIP). For each hnRNP C replicate, 4 µg hnRNP C1/C2 Antibody (4F4, Santa Cruz Biochnology #sc-32308) and 20 µL Dynabeads Protein G for Immunoprecipitation (ThermoFisher, #10003D) were coupled for 1 hour at room temperature before adding 600 µg of clarified HEK293T lysate and immunopurifying at 4° for 45-60 minutes. For FBF, two replicates of 4 mg clarified lysate were combined with 20 µL Fibrillarin Antibody (Bethyl, #A303-891A) and 20 µL Protein G Dynabeads; immunopurification was at 4° for 1 hour. The easyCLIP assay was performed as described in File S1.

### easyCLIP: library creation

The full easyCLIP protocol and all buffers are described in File S1. After harvesting, cells were thawed and lyzed with a microtip sonicator six times for five seconds each (10% power), with samples cooled by placement in dry ice between sonications. Lysates were then clarified by spinning at 14 krcf for 10 minutes at 4° and transferring the supernatant to a new tube. Concentrations were determined by BCA (see BCA section). To visualize protein expression levels, 15 µg of clarified lysates were used for western blotting. For immunopurification, typically 20 µL of anti-HA beads per sample were washed with NT2 buffer, then CLIP lysis buffer. Samples were diluted to 1-4 mg/mL during immunopurification, typically ~2 mg/ml. Immunopurification was 40 minutes to 1 hour at 4°. Samples were then washed once with H. Str. Buffer (10 minutes), H. Salt buffer (10 minutes), low salt buffer, and finally NT2 buffer, each with 1 mL. Samples were then stepped down with another wash to ~200 µL NT2 buffer. RNAse digestion was performed by diluting 2 µL 100 U/µL RNAse ONE to 1 U/µL in NT2 buffer, then diluting this to 0.025 U/µL in NT2 buffer with 16% PEG and adding 60 µL of this to each sample. The digestion was performed for 8-12 minutes at 30° with intermittent shaking. The digestion mixture was removed from the beads and 1 mL H. Str. Buffer was added. Samples were then washed twice with 1 mL NT2 buffer before being stepped down to ~200 µL NT2 buffer. Samples were then processed in the order (1) kinase, (2) 5’ ligation, (3) L5 barcodes combined, (4) phosphatase, (5) 3’ ligation, or in the order (1) phosphatase, (2) 3’ ligation, (3) L3 barcodes combined, (4) kinase, (5) 5’ ligation. Processing details and oligonucleotide sequences are in File S1. In either case, all samples were typically combined before being loaded into a single lane of a 4-12% NuPAGE Bis-Tris gel, run at 200V for ~45 minutes, and transferred to nitrocellulose at 400-500 mA for ~25 minutes. Membranes were then placed in PBS and immediately imaged in an Odyssey CLx machine. Membranes were cut using scalpels and put in 375 µL PK buffer with 25 µL Proteinase K and incubated for 40-60 minutes with shaking at 45-55°. In some cases, 2 µL of extracted RNA was then spotted on nylon and imaged. PK mixtures were added directly to 20 µL oligonucleotide(dT) beads and mixed at room temperature for 20 minutes. Alternatively, 2 M KCl was added and SDS was spun out, then 20 µL oligonucleotide(dT) beads were added and the samples were mixed at 4° for 20 minutes. Beads were washed once with biotin IP buffer, once with NT2 buffer, transferred to a PCR tube, then washed 3-4 times with PBS buffer. Samples were eluted in 14.4 water with 15 pmol reverse transcription primer by heating at 95° for 3 minutes and transferring to a new tube. Reverse transcription was performed by incubating for 40 minutes at 53° and 10 minutes at 55°, or in some cases for 40 minutes at 53° only. Reverse transcription product was then used directly for PCR as described in File S1.

### Ligation efficiency test by protein shift

The ligation efficiency test with hnRNP C was performed in three replicates. hnRNP C was purified by incubating 600 µg of clarified HEK293T lysate with 4 µg anti-hnRNP C1/C2 antibody for 1.5 hours at 4° as described previously^5^. Beads were RNAse digested and dephosphorylated as described previously, before being split 2:1. The split corresponding to 200 µg lysate was PNK phosphorylated and 5’ ligated as described in the easyCLIP protocol. The split corresponding to 400 µg was 3’ ligated as described previously, before being split in half. One 3’ ligated split was PNK phosphorylated and 5’ ligated as described in the easyCLIP protocol. All samples were then run on a 4-12% SDS-PAGE gel (NuPAGE), transferred to nitrocellulose and visualized as described previously. The amount of RNA that was neither 5’ nor 3’ ligated was determined by the following reasoning. First, let P5 be the probability of a 5’ ligation, and P3 be the probability of a 3’ ligation. Let a = RNA with no ligation; b = RNA with a 3’ ligation only; c = RNA with a 5’ ligation only; and d = RNA with a 5’ and 3’ ligation. Let T = the total amount of RNA. It follows that:

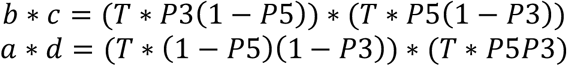

Rearranging terms shows that a*d = b*c. Since d, b, and c are determined by direct visualization of fluorescence, it follows that the RNA with no ligation (a) is also known.

### Fluorescence loss

20 µL of Streptavidin Dynabeads (ThermoFisher) per purification were washed three times with BIB, then combined with 2 µL of 5 µM biotin-anti-L5 RNA (10 pmol, ordered as /5BiosG/rUrArCrCrCrUrUrCrGrCrUrUrCrArCrArCrArCrArCrArArG from IDT, with an RNAse free HPLC purification). The oligonucleotide was captured for 20 minutes in 1 mL BIB, then washed with BIB, NT2, PBS (1X each) and resuspended in 50 µL BIB.

6.4 µL 2 M KCl was added to proteinase K-digested samples, and SDS was precipitated on ice for 15 minutes. SDS was spun out at 13 kRPM for 10 minutes. Dynabeads with 10 pmol biotin-anti-L5 RNA oligonucleotide in 50 µL BIB were then added to PK reactions and diluted to a total volume of 1 mL with BIB. The purification was carried out at 4° for 20 minutes. Beads were washed three times with BIB, twice with PBS, and eluted for 2 minutes at 95° in 15-20 µL water with 100 nM biotin.

10X NT2 was added to 1X final concentration, and PEG to 16% final concentration. 1 µL 100 U/µL RNAse ONE was added and samples incubated for 40 minutes at 37°. RNAse ONE was inactivated by adding 10% SDS to 0.1%. Shift buffer was added to 1X (25 mM Tris pH 7.5, 10 mM MgCl_2_, and 16% PEG400). 300-400 fmol labelled antisense oligos were added and samples were processed further as described for the ligation efficiency test by anti-sense oligo shift.

Shift oligos:

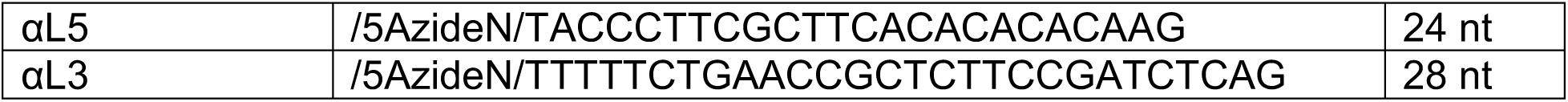

300-400 fmol labelled antisense oligonucleotides were added (max is ~500 fmol before signal cannot be quantified). The relative amount of shift oligonucleotide to input is important, as excessive oligonucleotide will create artifacts. Heat at 75° for 2 minutes, then let sample sit at room temperature for at least a minute. Create samples for two lanes of shift oligonucleotides at 300 fmol per lane (or however much was used to shift). Running the shift oligonucleotides at the same concentration used to shift is required to subtract background. Add 6X Ficoll/BPB buffer (15% Ficoll 400, 0.03% Bromophenol blue, 50 mM Tris pH 7.5) to 1X, but do not heat. For gel running buffer, add NaCl to 25 mM in 4° 0.5X TBE buffer. Samples were loaded on a 20% TBE gel and run gel 180V at 4° for one hour, replacing running buffer with 4° buffer every ~40 minutes. Finally, samples were transferred to nylon in 0.5X TBE buffer at 250 mA for 30 minutes.

### Ligation efficiency test by RNA shift

Samples of hnRNP C were prepared as normal for easyCLIP (File S1), and as described for the protein shift ligation efficiency test, up to the proteinase K extraction from nitrocellulose. To inactivate proteinase K, 6.4 µL 2M KCl per 400 µL of proteinase K extract was added, samples incubated at 4° for 15 minutes, and precipitated SDS removed by centrifugation at 13,000 RPM for 10 minutes at 4°.

Two sets of MyOne C1 Streptavidin beads were prepared, each using 13-20 µL MyOne C1 streptavidin beads per sample: one set for biotin purification and one for antisense oligonucleotide purification. Beads were washed three times with Biotin IP Buffer (BIB: 100 mM Tris pH 7.5, 1 M NaCl, 0.1% Tween-20, 1 mM EDTA). Those to be used for the biotin purification were then set aside until use. The set for anti-sense oligonucleotide purification were then incubated with 30 pmol anti-sense biotinylated oligonucleotide per µL resin in 1 mL BIB and rotated for 20 minutes at room temperature. Solution was removed and a second incubation with 15 pmol biotinylated oligonucleotide per µL resin was performed to ensure saturation. After incubation, anti-sense oligonucleotide beads were washed with BIB, NT2, PBS, and resuspended in 750 µL BIB. 50 µL of this bead solution was added to 400 µL BIB containing 20 nmol biotin and mixed. This solution was allowed to sit at room temperature for at least 5 minutes.

Proteinase K extract was bound to beads and incubated for 20 minutes at 4°. Supernatant was removed and beads were resuspended in 200 µL BIB, transferred to a PCR tube, rinsed with 200 µL NT2, washed with 200 µL PBS, and allowed to at least briefly reach 20-25°. After reaching room temperature, supernatant was removed and libraries eluted in 18 µL formamide at 65° for 2 minutes.

### Ligation efficiency test by anti-sense oligonucleotide shift

Beads were washed three times with BIB, twice with PBS, and eluted for 2 minutes at 95° in 15-20 µL water with 100 nM biotin. Add 10X NT2 to 1X, and PEG to 16% final concentration. Add 1 µL 100 U/µL RNAse ONE. Incubate 40 minutes at 37°. Add 10% SDS to 0.1% to inactivate RNAse ONE. Add shift buffer to 1X (25 mM Tris pH 7.5, 10 mM MgCl_2_, and 16% PEG400). Split the volume in three or four if doing separate shifts.

300-400 fmol labelled antisense oligos were added (max is 500 fmol before signal cannot be quantified). The relative amount of shift oligo to input is important, as excessive oligo will create artifacts. Samples were heated to 75° for 2 minutes, then cooled to room temperature at −0.1°/s. 6X Ficoll/BPB buffer (15% Ficoll 400, 0.03% Bromophenol blue, 50 mM Tris pH 7.5) was added to 1X before loading on a gel. For gel running buffer, NaCl to was added to 25 mM in 4° 0.5X TBE buffer. Samples were loaded on a 20% TBE gel and run at 180V at 4° for ~1-3 hours, replacing running buffer with 4° buffer every ~40 minutes. Finally, samples were transferred to nylon in 0.5X TBE buffer at 250 mA for 30 minutes.

### Generation of linear cDNA standards

Separately barcoded linear P3 and P6 fragments were ordered from IDT and stitched together by oligo extension. P3 fragments were of the following form, with X indicating the P3 barcode (Sequences in File 1). Fragments were mixed together in water and placed at room temperature before running the stitching reaction. Fragments were stitched together using Klenow fragment: 1 µL 100 µM of each oligo was combined with 10 µL NEBuffer 3.1 (10X), 1.5 µL of 2 mM dNTPs and 1 µL Klenow Fragment (exo-), in 100 µL reaction volumes. Reactions were incubated at 37° for 1 hour. 2 µL of Exonuclease I (NEB) was added to each reaction and incubated at 37° for 1 hour. Samples were purified with RNA clean and concentrator columns (Zymo) and eluted in 40 µL. Concentrations were determined by the dsDNA Qbit assay, and 1 µL of each sample was run on a 15% TBE-urea gel (NuPAGE). The dsDNA concentration obtained by Qbit was converted to a molar quantity using a molecular weight and fluorescence per fmol was determined by comparing the Qbit assay results and fluorescence on a TBE-urea gel. 3 fmol/µL samples were run again on a gel to determine concentration, diluted to 80 amol/µL, adjusted based on in-gel fluorescence, and finally diluted to 8 amol/µL. 0.3 µL of 8 amol/µL standards (2.4 amol) were added to CLIP PCR reactions. Consistency in final molar concentrations was evaluated by qPCR and adjusted towards the average.

### CLIP analysis: genomes

The GRCh38 genome Gencode release 29 and features were obtained from: ftp://ftp.ebi.ac.uk/pub/databases/gencode/Gencode_human/release_29/GRCh38.primary_assembly.genome.fa.gz ftp://ftp.ebi.ac.uk/pub/databases/gencode/Gencode_human/gencode.v29.primary_assembly.annotation.gtf.gz.

The STAR index was built using --sjdbOverhang 75. When assigning reads to genes after STAR mapping, only GTF features with transcript support level tsl1 or tslNA were included.

For repetitive elements, an alignment file from was downloaded from http://www.repeatmasker.org/. This was parsed to extract representatives, which were placed in an artificial chromosome separated by poly(N), and a gtf file for each representative was generated. A STAR index was built with --genomeSAindexNbases 5. The parameter genomeSAindexNbases must be set well below the default of 14 or building will be very slow. When mapping to the repeats chromosome, --alignIntronMax 1 was used to prevent the insertion of introns by STAR.

### CLIP analysis: read processing

Custom Python scripts (github.com/dfporter/easyCLIP) were used for all analysis. Raw fastq files were split by L5 and L3 barcodes allowing one nucleotide mismatches to the expected barcodes. Reads were first mapped to a custom-built chromosome of repetitive elements using STAR and “--alignEndsType EndToEnd”. Unmapped reads from this stage were then mapped to the regular genome using default parameters. Reads mapping the genome to remove multimapping reads and MAPQ < 10 reads. Mapping results from repetitive elements and the genome were combined, read mates removed, results converted to BED format, and PCR duplicates removed using the random hexamer UMI on the L5 adapter.

### CLIP analysis: read assignment

Reads were assigned to an RNA if they overlapped only that RNA, or if they overlapped a snoRNA element in any case. The strand was ignored for repetitive elements.

### CLIP analysis: statistics

Inputs to statistical analysis were either reads per million or reads per ten billion proteins, both treated the same. To speed up analysis, RNAs with a maximum count below five (reads per million reads, or per ten billion proteins) across all samples were dropped from all further analysis. For the randomly selected non-RBPs constituting background, if a replicate had no reads it was assigned one tenth the minimum positive count present in that dataset (i.e., if a dataset had one million reads, zeros were replaced with 0.1 reads per million). The average count across replicates for each protein was determined, resulting in a sample of eight values taken from the null distribution (one for each of the proteins CDK4, CHMP3, DCTN6, ETS2, IDE, ITPA, TPGS2 and UBA2). σ^2^/µ was essentially always above 2 for these samples, and were fit to a negative binomial using scipy^6^ and calculated P values accordingly before finally adjusting all P values for each protein by the Benjamini-Hochberg method into FDR equivalents.

### CLIP analysis: peak finding

For each RNA, reads spanning the genomic locus were converted into an array with the length of the genomic locus and each value representing the count of 5’ read ends mapping to that position. The values were smoothed by convolution using a box with length 50 for loci of at least 2,000 nucleotides, length 20 for 20-2,000 nucleotides, and length 10 for <200 nucleotides. If this array had a single maximum, it was taken to be the peak location. If there were multiple maxima (equal heights) and no maxima had more than a two nucleotide gap from another maxima, the peak was taken as the average position between the first and last maxima. If any maximum was more than two nucleotides from another maximum, the RNA was considered to have no peak.

